# Identification of a novel cobamide remodeling enzyme in the beneficial human gut bacterium *Akkermansia muciniphila*

**DOI:** 10.1101/2020.09.02.280537

**Authors:** Kenny C. Mok, Olga M. Sokolovskaya, Alexa M. Nicolas, Zachary F. Hallberg, Adam Deutschbauer, Hans K. Carlson, Michiko E. Taga

## Abstract

The beneficial human gut bacterium *Akkermansia muciniphila* provides metabolites to other members of the gut microbiota by breaking down host mucin, but most of its other metabolic functions have not been investigated. *A. muciniphila* is known to use cobamides, the vitamin B_12_ family of cofactors with structural diversity in the lower ligand, though the specific cobamides it can use have not been examined. We found that growth of *A. muciniphila* strain Muc^T^ was nearly identical with each of seven cobamides tested, in contrast to nearly all bacteria that have been studied. Unexpectedly, this promiscuity is due to cobamide remodeling – the removal and replacement of the lower ligand – despite the absence of the canonical remodeling enzyme CbiZ in *A. muciniphila*. We identified a novel enzyme, CbiR, that is capable of initiating the remodeling process by hydrolyzing the phosphoribosyl bond in the nucleotide loop of cobamides. CbiR does not share homology with other cobamide remodeling enzymes or B_12_-binding domains, and instead is a member of the AP endonuclease 2 enzyme superfamily. We speculate that CbiR enables bacteria to repurpose cobamides they otherwise cannot use in order to grow under a cobamide-requiring condition; this function was confirmed by heterologous expression of *cbiR* in *E. coli*. Homologs of CbiR are found in over 200 microbial taxa across 22 phyla, suggesting that many bacteria may use CbiR to gain access to the diverse cobamides present in their environment.

**Importance:** Cobamides, the vitamin B_12_ family of cobalt-containing cofactors, are required for metabolism in all domains of life, including most bacteria. Cobamides have structural variability in the lower ligand, and selectivity for particular cobamides has been observed in most organisms studied to date. Here, we discover that the beneficial human gut bacterium *Akkermansia muciniphila* can use a diverse range of cobamides due to its ability to change the cobamide structure via “cobamide remodeling”. We identify and characterize the novel enzyme CbiR that is necessary for initiating the cobamide remodeling process. The discovery of this enzyme has implications not only for understanding the ecological role of *A. muciniphila* in the gut, but for other bacteria that carry this enzyme as well.

## Introduction

The human gut microbiota is composed of diverse communities of microbes that play important roles in human health (1-4). Disruption of the composition of the microbiota, known as dysbiosis, is associated with numerous disease states (5-9). While the immense complexity and interindividual variability of the microbiota have made it challenging to identify the specific functions of most community members, particular taxa are starting to be linked to health and disease (10-12), with the bacterium *Akkermansia muciniphila* recently emerging as a beneficial microbe due to its distinctive metabolic capabilities (13, 14).

*A. muciniphila* is thought to benefit the host by inducing mucus production, improving gut barrier function, and stimulating a positive inflammatory response (15-24). *A. muciniphila* is one of few bacteria capable of using mucin, the main component of mucus, as a sole carbon, nitrogen, and energy source (25). Mucin degradation products released by *A. muciniphila* are used as carbon sources by butyrate-producing bacteria and likely other bacteria, and for this reason *A. muciniphila* is thought to be a keystone species in the gut (26, 27). In addition to providing metabolites to neighboring microbes, in coculture *A. muciniphila* can use a cobamide cofactor, pseudocobalamin (pCbl, Fig. 1A), provided by *Eubacterium hallii* for the production of propionate (26). Both butyrate and propionate positively affect host metabolism and immune function (28-30).

**Figure 1.**
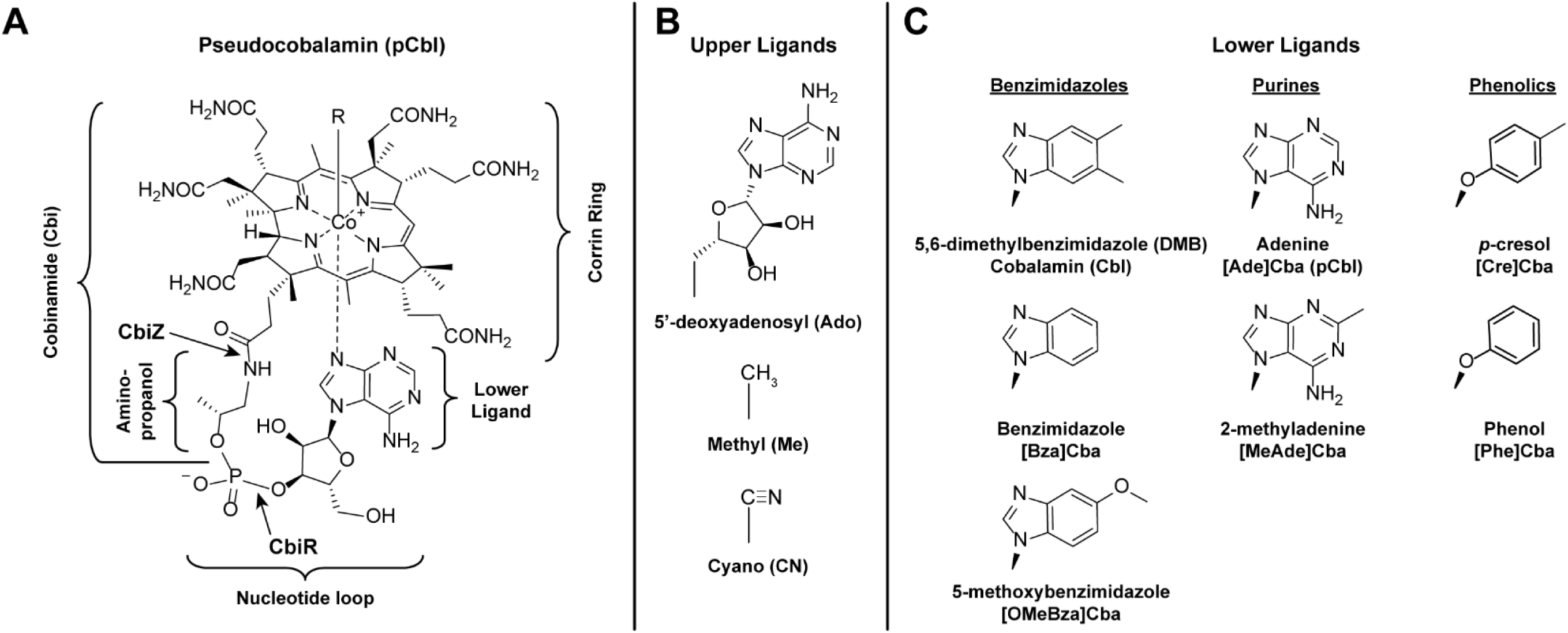
Cobamide structures. A. Structure of pCbl. All cobamides are composed of a corrin ring containing a central cobalt ion and an upper (R) and lower ligand. In pCbl, the lower ligand is adenine. The lower ligand, together with the ribose and phosphate moieties, comprise the nucleotide loop, which is covalently attached to the corrin ring via an aminopropanol linker. The bonds hydrolyzed by the CbiZ amidohydrolase and the CbiR phosphodiesterase are indicated with arrows. The part of the molecule comprising Cbi is shown. Cobamides and their corrin-containing biosynthetic precursors and degradation products are together known as corrinoids. B. Upper ligands (R) in cobamides, the catalytic center of the cofactor; prefixes used in the text to denote the upper ligand are shown in parentheses. C. The three chemical classes of lower ligands present in cobamides. The structures of the seven cobamide lower ligands used in this study are shown. Names of the lower ligand base and abbreviations used for the corresponding cobamides are given below the structures.

Cobamides are a family of cobalt-containing corrinoid cofactors that include vitamin B_12_ (cobalamin, Cbl), an essential micronutrient for humans. Cobamides are required by organisms in all domains of life, but are synthesized only by a subset of prokaryotes (31-33). While some strains of *A. muciniphila* were shown or predicted to produce cobamides *de novo*, the type strain, Muc^T^, is incapable of *de novo* cobamide production (34). Instead, strain Muc^T^ and most other *A. muciniphila* strains are predicted to be capable of cobinamide (Cbi, Fig. 1A) salvaging (33, 34), a process in which a cobamide is synthesized from the late precursor Cbi (35). Thus, the four cobamide-dependent metabolic pathways present in *A. muciniphila* function in most strains, including Muc^T^, only when a cobamide or a late precursor such as Cbi is provided by another organism. Several other human gut bacteria have similarly been found to use cobamide cofactors but are unable to produce them *de novo*, including *Bacteroides fragilis, Bacteroides thetaiotaomicron, Bacteroides vulgatus, Clostridioides difficile, Enterococcus faecalis, Escherichia coli*, and *Parabacteroides distasonis* (36-40). In addition to these specific examples, genomic analysis suggests that dependence on cobamide-producing microbes is widespread in the gut and other environments: 58% of human gut bacteria and 49% of all sequenced bacteria are predicted to use cobamides but lack the capacity to produce them (33).

A feature that sets cobamides apart from other enzyme cofactors is that different microbes produce structurally distinct cobamides (41). This variability is mostly limited to the lower ligand, which can be benzimidazolyl, purinyl, or phenolyl bases (Fig. 1C). Individual cobamide-producing bacteria typically synthesize only one type of cobamide, but microbial communities have been found to contain four to eight different cobamides or cobamide precursors (42-45). A study of 20 human subjects showed that the human gut is dominated by the purinyl class of cobamides, with benzimidazolyl and phenolyl cobamides and Cbi also present (42). The structural diversity in cobamides impacts growth and metabolism, as most organisms studied to date are selective in their cobamide use (39, 46-54). For example, the human gut bacterium *B. thetaiotaomicron* can use benzimidazolyl and purinyl, but not phenolyl, cobamides (37); *Dehalococcoides mccartyi* and most eukaryotic algae are selective for particular benzimidazolyl cobamides (55-57); and *Sporomusa ovata* requires phenolyl cobamides (58). Thus, microbes that depend on cobamides produced by others may struggle to grow in environments lacking their preferred cobamides. However, some organisms have evolved mechanisms of acquiring the specific cobamides that function in their metabolism. For example, bacterial cobamide uptake can be somewhat selective, as shown in a study in *B. thetaiotaomicron* (37). Another strategy used by some microbes is cobamide remodeling, the removal and replacement of the lower ligand.

Cobamide remodeling was first described in *Rhodobacter sphaeroides* (59), but has also been observed in the bacteria *D. mccartyi* and *Vibrio cholerae* and the algae *Pavlova lutheri* and *Chlamydomonas reinhardtii* (45, 48, 55, 57). In each case, cobamide remodeling enables the organism to repurpose a cobamide that poorly supports growth. In *R. sphaeroides*, the cobamide remodeling process is initiated by the enzyme CbiZ, which hydrolyzes the amide bond adjacent to the aminopropanol linker (Fig. 1A) (59); in subsequent steps, cobamide biosynthesis is completed with a different lower ligand via the activity of six gene products, most of which are also required for Cbi salvaging. *In vitro, R. sphaeroides* CbiZ hydrolyzes pCbl but not Cbl (59). This specificity is thought to drive the conversion of pCbl, a cofactor that *R. sphaeroides* cannot use, into Cbl, which functions in its metabolism. *D. mccartyi* also has homologs of *cbiZ* (57), while cobamide remodeling in *V. cholerae* was recently shown to involve the cobamide biosynthesis enzyme CobS (48). The genes required for cobamide remodeling in algae have not been identified. Nevertheless, *A. muciniphila* does not encode a homolog of *cbiZ*, and therefore we assessed the cobamide selectivity of *A. muciniphila* strain Muc^T^ to understand the cobamide metabolism of this beneficial gut bacterium.

Here we show that *A. muciniphila* strain Muc^T^ is able to grow equivalently when provided a variety of cobamides. We found that this lack of selectivity is due to the unexpected ability of *A. muciniphila* to remodel cobamides. We identified a previously uncharacterized phosphodiesterase in *A. muciniphila* that we named CbiR, which initiates the remodeling process by hydrolyzing cobamides. Heterologous expression in *E. coli* shows that CbiR expands access to a cobamide that does not otherwise support growth. Homologs of CbiR are present in the genomes of microbes in diverse habitats from 22 phyla, and phylogenetic analysis establishes CbiR as a new, distinct clade within the AP endonuclease 2 superfamily. These observations enhance the understanding of the metabolic roles of *A. muciniphila* and improve our ability to predict cobamide-dependent physiology in other bacteria.

## Results

### *A. muciniphila* strain Muc^T^ salvages Cbi to produce pCbl

*A. muciniphila* strain Muc^T^ lacks most of the genes required for cobamide synthesis and does not produce cobamides *de novo* (34), but it is predicted to be capable of Cbi salvaging (33). To test this prediction, we extracted corrinoids from *A. muciniphila* cultured with and without Cbi and analyzed the corrinoid composition of the samples by high-performance liquid chromatography (HPLC). When cultured without Cbi, no corrinoids were detected in the extractions (Fig. 2A). However, when Cbi was added to the growth medium, a cobamide with the same retention time and nearly identical UV-Vis spectrum to pCbl was detected by HPLC (Fig. 2A). Mass spectrometry (MS) analysis corroborated that this cobamide is pCbl (Fig. S1).

**Figure 2.**
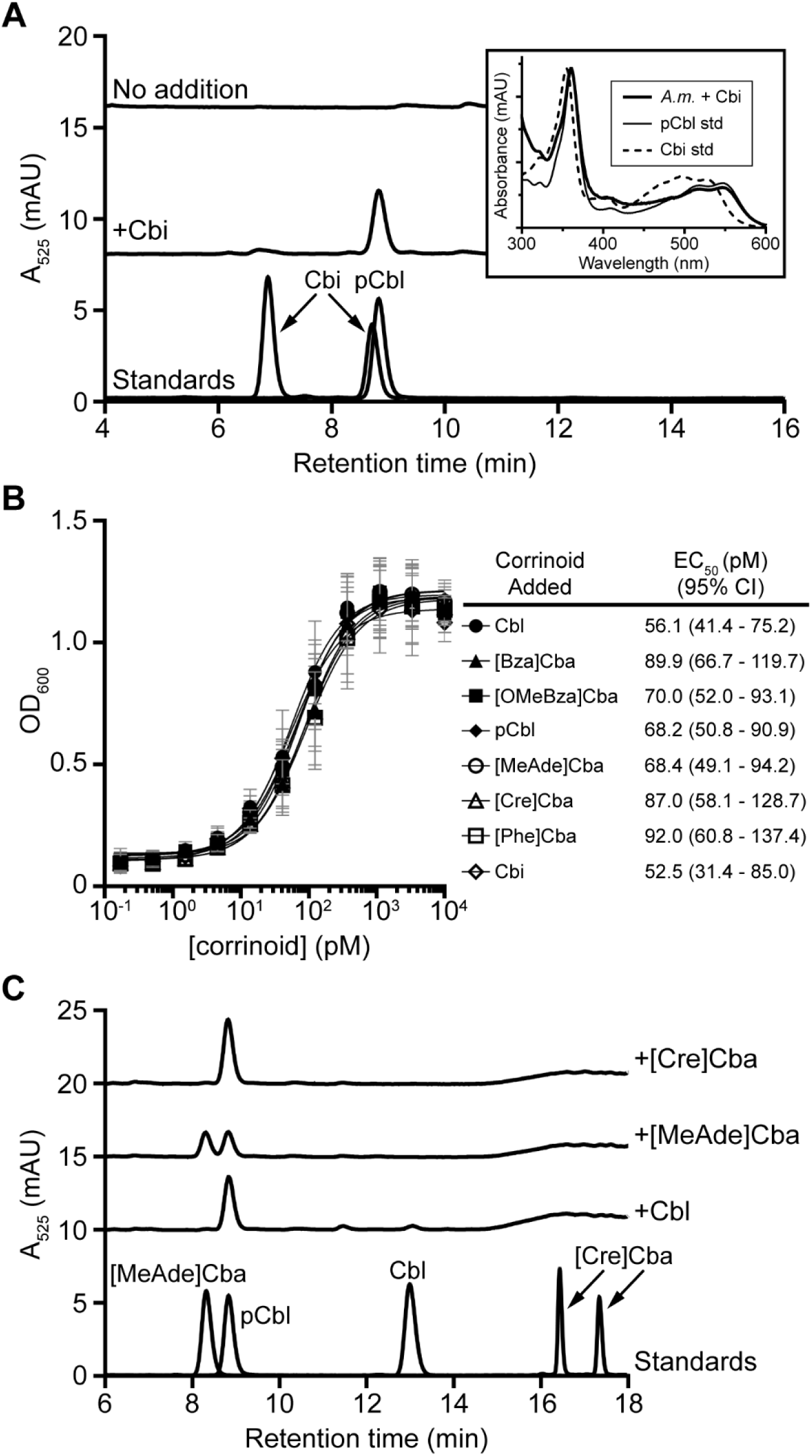
*A. muciniphila* strain Muc^T^ can salvage Cbi and remodel cobamides. A. HPLC analysis of corrinoid extractions from *A. muciniphila* grown with or without 10 nM Cbi for 72 h shows that *A. muciniphila* can salvage Cbi. Standards for Cbi and pCbl are shown at the bottom. Comparison of UV-Vis spectra (inset) of the HPLC peaks at 8.8 min shows that the corrinoid produced by *A. muciniphila* (*A*.*m*.) grown with Cbi (thick line) is similar to a pCbl standard (std) (thin line) and not a Cbi standard (dashed line). Spectra were normalized to each other based on their maxima to aid comparison. B. *A. muciniphila* growth under methionine-deplete conditions. OD_600_ of saturated cultures is shown after 29 h of growth with the indicated concentrations of each corrinoid. EC_50_ values and their 95% confidence intervals for each corrinoid are given in the table. Data points and error bars represent the mean and standard deviation, respectively, of three biological replicates. The results are representative of three independent experiments. C. HPLC analysis of corrinoid extractions from *A. muciniphila* grown with 10 nM Cbl, [MeAde]Cba, or [Cre]Cba for 72 h shows that *A. muciniphila* remodels cobamides to pCbl. Cobamide standards are shown at the bottom.

### *A. muciniphila* strain Muc^T^ does not show cobamide selectivity

Having established that *A. muciniphila* strain Muc^T^ cannot synthesize cobamides without the addition of a precursor, we next examined which cobamides it is capable of using by measuring growth in the presence of various cobamides under a cobamide-requiring condition. A homolog of the cobamide-dependent methionine synthase MetH is encoded in the genome of *A. muciniphila*. Because *A. muciniphila* lacks a homolog of the cobamide-independent methionine synthase MetE, growth in methionine-deplete medium is expected to require cobamide addition. We found this to be the case, as the addition of Cbi or any of the seven cobamides tested was necessary to support growth of *A. muciniphila* (Fig. 2B). Surprisingly, however, *A. muciniphila* shows essentially no cobamide selectivity, with less than twofold variation in the cobamide concentrations resulting in half-maximal growth (EC_50_) (Fig. 2B).

### *A. muciniphila* strain Muc^T^ remodels cobamides to pCbl

The ability of all of the tested cobamides to support nearly identical growth of *A. muciniphila* could be due to promiscuity in its cobamide-dependent methionine synthase. Alternatively, *A. muciniphila* could remodel cobamides, despite the absence of a homolog of *cbiZ* in its genome. If cobamide remodeling occurs in *A. muciniphila*, exogenously supplied cobamides will be altered by the bacterium. Therefore, we extracted corrinoids from *A. muciniphila* cultures supplemented with Cbl, [Cre]Cba or [MeAde]Cba to determine whether the added cobamides could be recovered. HPLC analysis revealed that none of the Cbl or [Cre]Cba, and only half of the [MeAde]Cba, remained in the extractions. This loss of the added cobamide coincided with the appearance of a new cobamide that co-eluted with pCbl (Fig. 2C). MS analysis confirmed that this cobamide is indeed pCbl (Fig. S2). These results demonstrate that *A. muciniphila* remodels cobamides to pCbl.

### Identification and characterization of a novel cobamide remodeling enzyme in *A. muciniphila*

The identification of cobamide remodeling activity despite the absence of a *cbiZ* homolog in the genome suggested that a novel enzyme capable of hydrolyzing cobamides is present in *A. muciniphila*. We reasoned that the gene encoding this enzyme could be located near the cobamide biosynthesis and salvaging genes *cobDQ, cbiB, cobT, cobS*, and *cobU*, some or all of which would be required for completion of the remodeling process. These five genes are found at a single locus in the *A. muciniphila* genome that also contains an ORF with unknown function, annotated as Amuc_1679 (Fig. 3A). Amuc_1679 is predicted to encode a protein with a conserved (β/α)_8_ TIM barrel domain from the AP endonuclease 2 superfamily (pfam01261). This superfamily is composed of several enzymes including endonuclease IV, which hydrolyzes phosphodiester bonds at apurinic or apyrimidinic (AP) sites in DNA (60). The proximity of Amuc_1679 to genes involved in cobamide biosynthesis and the presence of a phosphodiester bond connecting the lower ligand to the aminopropanol linker suggested that Amuc_1679 might play a role in cobamide biology in *A. muciniphila*. Further, homologs of this gene in other bacteria are also found in loci containing similar cobamide biosynthesis enzymes (Fig. 3A, Fig. S3).

**Figure 3.**
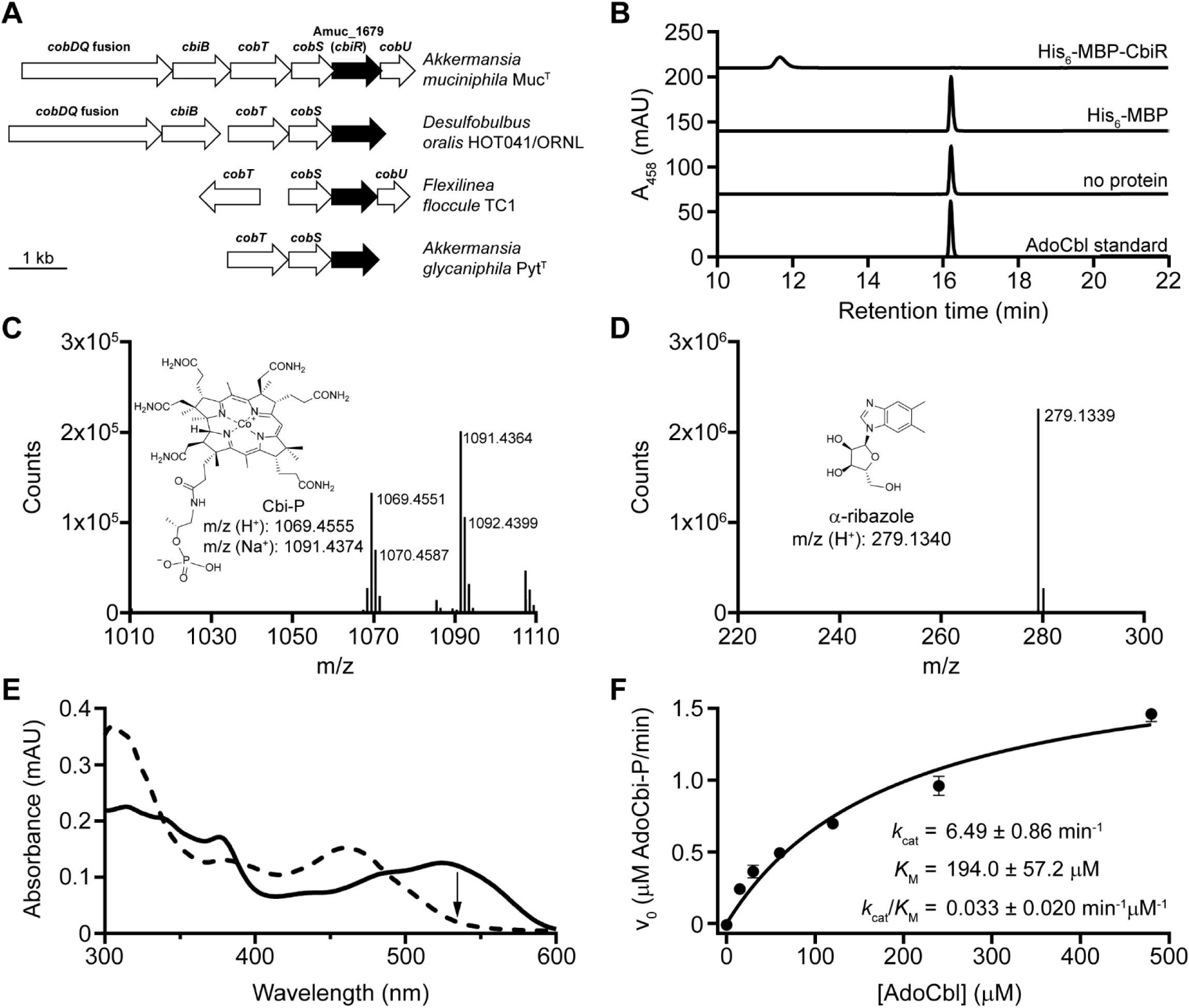
Purified CbiR hydrolyzes AdoCbl to form AdoCbi-P and α-ribazole *in vitro*. A. *A. muciniphila* Amuc_1679 (*cbiR*) and homologs in other bacteria (black arrows) are located near cobamide biosynthesis genes (white arrows). An expanded genomic comparison is shown in Figure S3. B. Purified CbiR converts AdoCbl to another corrinoid compound *in vitro* anaerobically. HPLC analysis of reactions containing 10 µM AdoCbl incubated for 4 h with 0.1 µM His_6_-MBP-CbiR, 0.1 µM His_6_-MBP, or no protein is shown. An AdoCbl standard is shown at bottom. C. The corrinoid product of His_6_-MBP-CbiR was purified by HPLC, exposed to light to remove the adenosyl upper ligand, and analyzed by MS. The structure and predicted m/z for Cbi-P are shown for comparison. D. The second product of the *in vitro* reaction with His_6_-MBP-CbiR and AdoCbl was purified by HPLC and analyzed by MS. The structure and predicted m/z for α-ribazole are shown for comparison. E. Comparison of the UV-Vis spectra before (solid line) and after completion (dashed line) of the reaction of His_6_-MBP-CbiR with 30 µM AdoCbl shows a decrease in absorbance at 534 nm (arrow). F. Michaelis-Menten kinetic analysis of His_6_-MBP-CbiR with AdoCbl. Reactions contained 0.3 µM His_6_-MBP-CbiR. Points and error bars represent the mean and standard deviation, respectively. Kinetic constants were determined from two independent experiments, each with three technical replicates.

To determine whether Amuc_1679 encodes an enzyme that can hydrolyze cobamides, we overexpressed and purified Amuc_1679 with N-terminal hexahistidine (His_6_) and maltose-binding protein (MBP) tags for analysis of its activity *in vitro* (Fig. S4A). First, we tested whether a new product was formed when the protein was incubated with coenzyme B_12_ (AdoCbl), an active cofactor form of Cbl. We observed complete conversion of AdoCbl to a new corrinoid compound in reactions performed under anaerobic conditions (Fig. 3B). MS analysis showed that two reaction products, cobinamide-phosphate (Cbi-P) and α-ribazole, were formed, indicating hydrolysis of the phosphoribosyl bond of AdoCbl (Fig. 3C, D). Notably, Amuc_1679 targets a bond distinct from the enzyme CbiZ (Fig. 1A) (59). In keeping with the tradition of naming cobamide biosynthesis and remodeling enzymes with the “Cbi” prefix, we henceforth refer to Amuc_1679 as CbiR.

We were able to monitor CbiR activity continuously by measuring the rate of decrease in absorbance at 534 nm (A_534_), as the reaction is characterized by a change in the UV-Vis spectrum that reflects the loss of AdoCbl and formation of AdoCbi-P (Fig. 3E). With this method, we found that the reaction proceeded only in the absence of oxygen, and additionally that the reaction requires the reducing agent DTT and is inhibited by the metal chelator EDTA (Fig. S4B). Using the same method, we determined the reaction kinetics of His_6_-MBP-CbiR under steady-state conditions at a range of AdoCbl concentrations (Fig. 3F). Based on a fit to the Michaelis-Menten model, the reaction of His_6_-MBP-CbiR with AdoCbl exhibited a *K*_M_ and *k*_cat_ for AdoCbl of 194 μM and 6.5 min^-1^, respectively. Similarly, His_6_-MBP-CbiR hydrolyzes MeCbl, the active cofactor form used by MetH and other methyltransferases, to MeCbi-P (Fig. S4C), with comparable kinetic parameters (Fig. S4D), indicating that AdoCbl and MeCbl are equally suitable substrates for His_6_-MBP-CbiR.

### *A. muciniphila* CbiR can hydrolyze several different cobamides *in vitro*

To determine the substrate selectivity of CbiR, His_6_-MBP-CbiR activity was measured *in vitro* with seven different cobamides. Adenosylated cobamides with purinyl or phenolyl lower ligands do not show UV-Vis spectra distinct from AdoCbi-P under the reaction conditions, and thus activity was measured by HPLC. Each cobamide was completely converted to AdoCbi-P following an 18 h incubation, demonstrating that all of the cobamides are substrates for CbiR (Fig. 4A). The specific activities of His_6_-MBP-CbiR with each cobamide are similar, with 4-fold differences among the benzimidazolyl and purinyl cobamides and slightly higher activity with phenolyl cobamides (Fig. 4B). These specific activities are similar to, though slightly lower than that previously reported for CbiZ with Ado-pCbl (70 nmol/mg/min, (59)), albeit under somewhat different reaction conditions.

**Figure 4.**
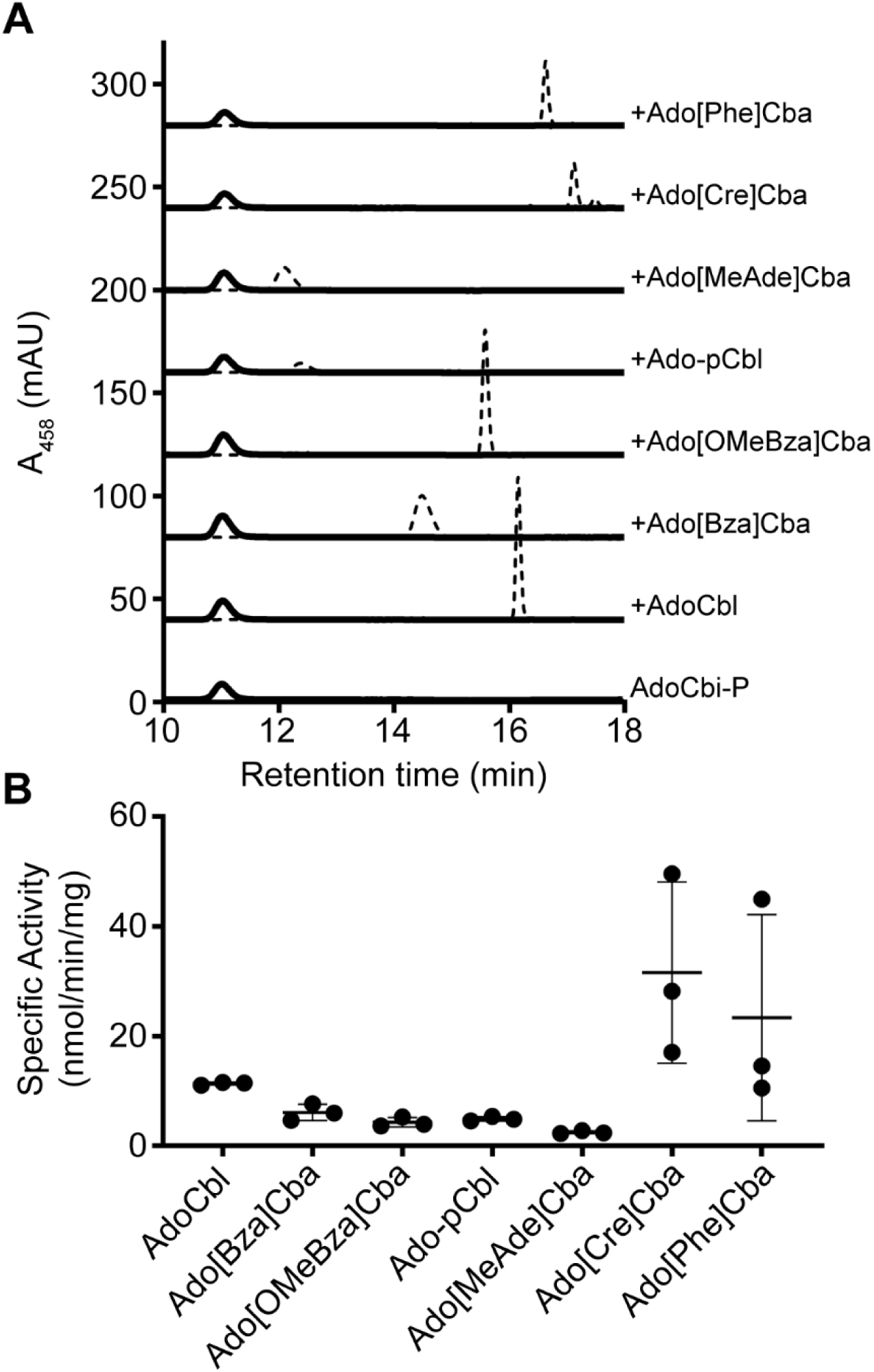
CbiR hydrolyzes many cobamides to form AdoCbi-P. A. HPLC analysis of *in vitro* reactions with different cobamides (10 µM), quenched after 18 h, is shown for reactions containing 0.1 µM His_6_-MBP-CbiR (solid lines) or without enzyme (dashed lines). A sample of purified AdoCbi-P is shown at the bottom. B. Specific activity of His_6_-MBP-CbiR with different cobamide substrates. 0.3 µM His_6_-MBP-CbiR was incubated with 30 µM of each cobamide individually and the rate of AdoCbi-P production was determined based on HPLC measurements at three time points. The lines represent the mean and standard deviation for three independent experiments.

### Expression of *cbiR* in *E. coli* enables expanded cobamide use

Given that the product AdoCbi-P can be used as a precursor for construction of a different cobamide, we hypothesize that CbiR activity enables bacteria to remodel cobamides, and therefore to gain access to cobamides in the environment that they otherwise may not be able to use. Because methods for targeted inactivation of genes in *A. muciniphila* have not been established, we used engineered *E. coli* strains to test this hypothesis. Like *A. muciniphila* strain Muc^T^, *E. coli* MG1655 cannot synthesize cobamides *de novo*, but its genome has the cobamide biosynthesis genes *cobT, cobS, cobU*, and *cobC* that should allow *E. coli* to convert AdoCbi-P into a cobamide (61). We first tested whether *A. muciniphila* CbiR is functional in *E. coli*. Indeed, expression of *cbiR* on a plasmid in a Δ*cobTSU* Δ*cobC* background results in the loss of added Cbl, pCbl, [MeAde]Cba, and [Cre]Cba and the formation of two new corrinoid compounds (Fig. 5A). One of the products co-elutes with AdoCbi-P (Fig. 5A), and MS analysis confirmed that the dominant ion matches the m/z expected for Cbi-P (Fig. S5A). The second product has an m/z consistent with Cbi (Fig. S5B), which likely forms intracellularly by hydrolysis of the phosphate group of AdoCbi-P. Neither product was detected in an *E. coli* strain containing the empty vector (Fig. 5A, dashed lines). Therefore, the activity of CbiR that we observed *in vitro* can be recapitulated in aerobically cultured *E. coli*.

**Figure 5.**
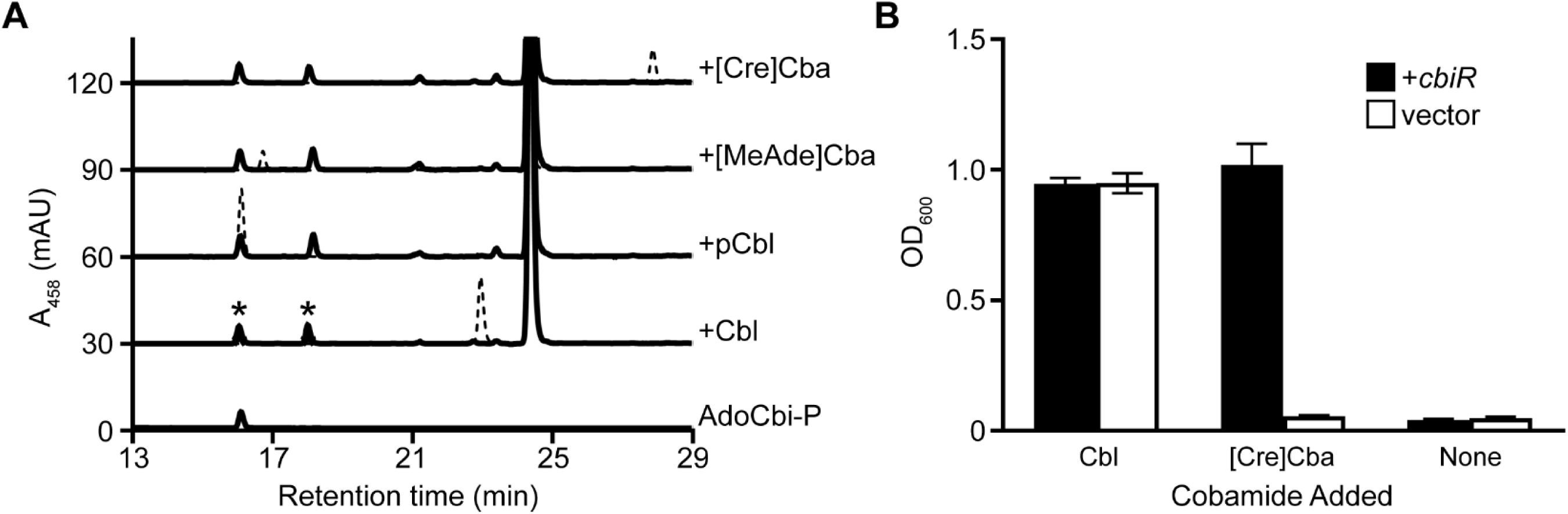
CbiR mediates cobamide remodeling in *E. coli*. A. CbiR hydrolyzes cobamides in *E. coli. A. muciniphila cbiR* was expressed in an *E. coli* strain with the *cobTSU* operon and *cobC* gene deleted to prevent modification of cobamide hydrolysis products. Corrinoid extractions of *E. coli* strains carrying pETmini-*cbiR* (solid lines) or the pETmini empty vector (dashed lines), grown with 10 nM Cbl, pCbl, [MeAde]Cba, or [Cre]Cba were analyzed by HPLC. A sample of purified AdoCbi-P is shown at the bottom. Corrinoids labeled with asterisks were purified for MS analysis (Fig. S5). The large peak at 24.5 min corresponds to a flavin that is present in all of the corrinoid extractions. B. Expression of *A. muciniphila cbiR* enables growth of *E. coli* on ethanolamine. Wild type *E. coli* MG1655 harboring pETmini-*cbiR* (black bars) or the pETmini empty vector (white bars) was cultured in minimal medium containing ethanolamine as the sole nitrogen source and 1 µM DMB. Cultures were supplemented with 100 nM Cbl, [Cre]Cba, or no cobamide and OD_600_ measurements are shown after 72 h of growth. Bars and error bars represent the mean and standard deviation, respectively, of three biological replicates.

Catabolism of ethanolamine in *E. coli* requires the cobamide-dependent enzyme ethanolamine ammonia lyase, which is capable of using Cbl as a cofactor but is not functional with [Cre]Cba (62). We took advantage of this selectivity to design a cobamide remodeling-dependent growth assay in *E. coli*. In minimal medium supplemented with [Cre]Cba and 5,6-dimethylbenzimidazole (DMB, the lower ligand of Cbl), with ethanolamine as the sole nitrogen source, *E. coli* should be able to grow only if it can remodel [Cre]Cba to Cbl. Cbl, as expected, promotes growth of *E. coli* under this condition regardless of whether *cbiR* is present (Fig. 5B). In contrast, when [Cre]Cba is added, growth is observed only in the strain expressing *cbiR*, suggesting that CbiR activity enables *E. coli* to convert [Cre]Cba into Cbl (Fig. 5B). A cobamide with a retention time and m/z matching that of Cbl was detected in a corrinoid extraction of *E. coli* grown with [Cre]Cba and DMB when expressing CbiR, confirming that cobamide remodeling to Cbl occurred (Fig. S6). These results demonstrate that expression of CbiR expands the range of cobamides accessible to *E. coli*, and suggests that cobamide remodeling may serve a similar purpose in *A. muciniphila*.

### CbiR is a member of the AP endonuclease 2 superfamily and is found in diverse bacteria

Analysis of the sequence of CbiR revealed that it is not similar to CbiZ or *V. cholerae* CobS, the other enzymes known to have cobamide remodeling activity. Instead, CbiR is a member of the AP endonuclease 2 superfamily, which includes the enzymes endonuclease IV, 2-keto-myo-inositol dehydratase, xylose isomerase, and other sugar isomerases and epimerases. A phylogenetic tree of this superfamily shows that the CbiR homologs identified by a BLAST search that are encoded in genomic loci containing cobamide biosynthesis genes form a single, distinct clade within the superfamily (Fig. 6A). Some of the biochemically characterized enzymes in the superfamily require metal cofactors for activity, and between one and three metal ions are found in nearly all X-ray crystal structures of enzymes from the superfamily (63-77). A metal cofactor may also be required for CbiR function; in addition to the inhibition by the metal chelator EDTA (Fig. S4B), CbiR homologs contain conserved His, Asp, and Glu residues that, in the characterized members of the superfamily, are involved in metal coordination (Fig. 6B). Furthermore, single mutations in many of these conserved residues in CbiR eliminated most or all of its AdoCbl hydrolysis activity when expressed in *E. coli* (Fig. 6C). These results demonstrate that CbiR shares both sequence and functional features common to the AP endonuclease 2 superfamily, and represents a new function within the superfamily.

**Figure 6.**
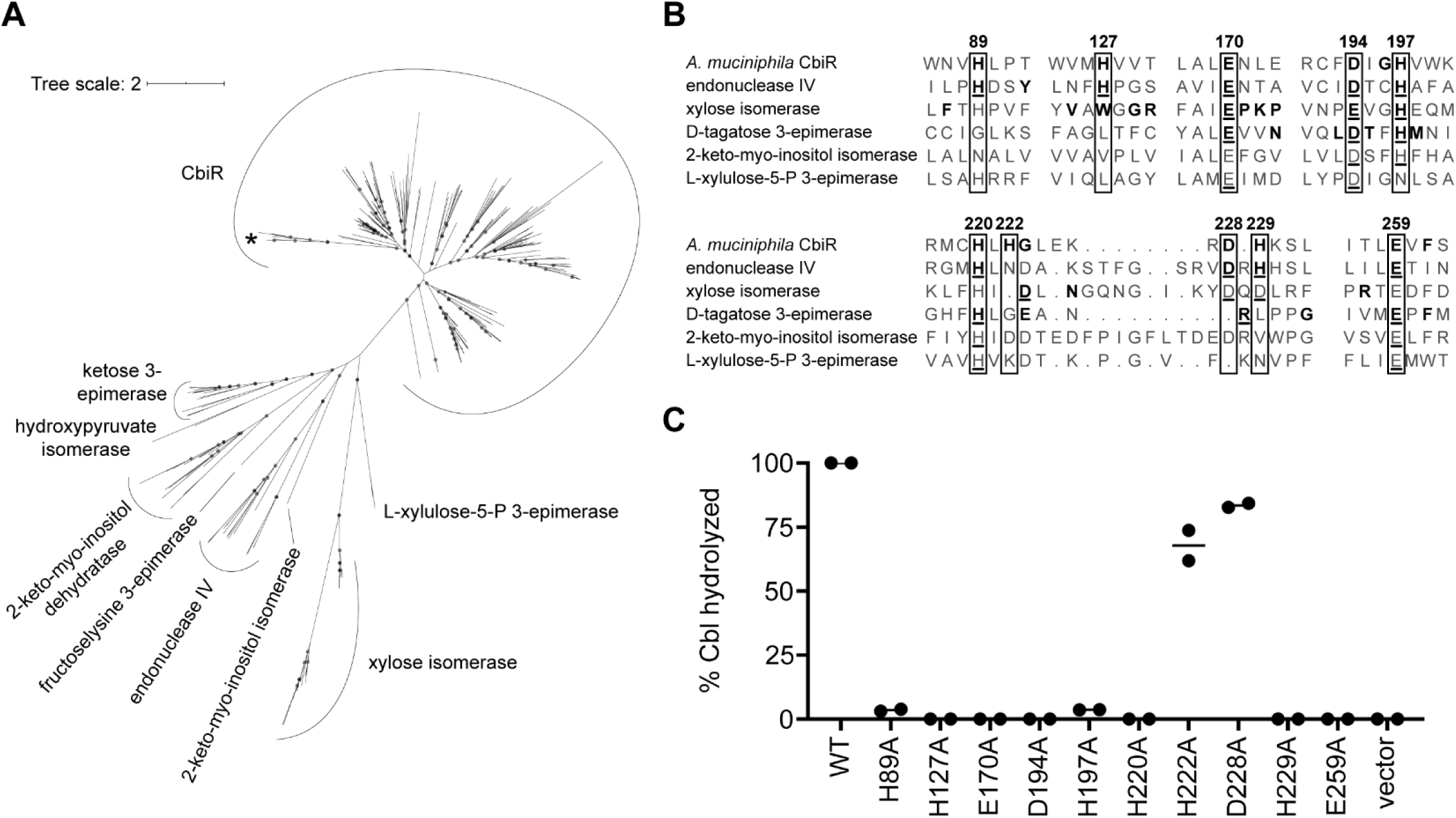
CbiR is a newly described member of the AP endonuclease 2 superfamily. A. Maximum likelihood tree of CbiR homologs and members of the AP endonuclease 2 superfamily that have been experimentally characterized. *A. muciniphila* CbiR is indicated by an asterisk. CbiR homologs included in the tree were identified in a BLAST search queried with *A. muciniphila* CbiR with E values lower than 10^−14^ and encoded adjacent to and in the same orientation as one or more cobamide biosynthesis gene (Table S1). Superfamily member sequences are listed in Table S2. Gray circles overlaid on tree nodes represent bootstrap values of >95%. The scale bar corresponds to the average number of substitutions per site across the alignment. B. Sequence alignment of regions containing highly conserved His, Asp, and Glu residues in *A. muciniphila* CbiR and representative sequences of biochemically and structurally characterized enzyme classes in the AP endonuclease 2 superfamily (*E. coli* endonuclease IV, *Streptomyces rubiginosus* xylose isomerase, *Pseudomonas cichorii* D-tagatose 3-epimerase, *Bacillus subtilis* 2-keto-myo-inositol isomerase, *E. coli* L-xylulose-5-P 3-epimerase). Numbers correspond to positions in *A. muciniphila* CbiR. For CbiR, endonuclease IV, xylose isomerase, and D-tagatose (ketose) 3-epimerase, bolded residues represent conserved amino acids in the enzyme classes. Underlined residues indicate amino acids in the X-ray crystal structures that interact with the metal cofactor(s), and with the substrate in the case of *P. cichorii* D-tagatose 3-epimerase. C. Mutational analysis of *A. muciniphila* CbiR. Corrinoids were extracted from cultures of *E. coli* Δ*cobTSU* Δ*cobC* strains carrying pETmini-*cbiR* (WT), pETmini-*cbiR* with the specified alanine mutations, or the pETmini empty vector grown for 20 h with 75 nM Cbl and analyzed by HPLC. The y-axis represents the combined amount of AdoCbi-P and AdoCbi present out of the total adenosylated corrinoids extracted. Minimal amounts of CNCbl were detected in the mutant extractions and were excluded from the analysis. The total intracellular corrinoid content was similar between samples except for WT and D228A, which had 4- and 2.5-fold higher levels, respectively. Lines show the mean of the two independent experiments.

Finally, we investigated the prevalence of CbiR across sequenced organisms by examining genomes with *cbiR* homologs. The *cbiR* gene commonly occurs in the *Akkermansia* genus, as a search of the 191 available genomes in the NCBI database found that 184 have *cbiR*. Additionally, 282 homologs of *A. muciniphila* CbiR with Expect values below 10^−3^ were identified by BLAST in the genomes of 275 bacterial and 1 archaeal taxa from diverse habitats including aquatic environments, sewage, digesters, oil spills, bioreactors, soil, and human and animal hosts (Table S1). While 76% are found in the phyla Chlorobi, Chloroflexi, and Proteobacteria, CbiR homologs are also found in 19 other phyla including six candidate phyla and two candidate divisions (Fig. S3, Table S1), with relatively few in the PVC superphylum, to which *A. muciniphila* belongs. Similar to *A. muciniphila*, 80% of the *cbiR* homologs are located adjacent to genes involved in cobamide biosynthesis (Table S1), suggesting that they, too, function in cobamide remodeling. It is therefore likely that cobamide remodeling initiated by CbiR occurs in diverse bacteria and environments.

## Discussion

Cobamides are considered to be important modulators of mammalian gut ecosystems because they are involved in several metabolic pathways, their production is limited to a subset of prokaryotes, and their diverse structures are differentially accessible to different microbes (54, 78, 79). *A. muciniphila* has been shown to have positive effects on host metabolism, gut barrier function, and the inflammatory response (15-24), yet knowledge of its metabolic and ecological roles in the gut remains incomplete. Previous studies showed that *A. muciniphila* strain Muc^T^ is unable to produce cobamides *de novo* (34), but can use pCbl produced by *E. hallii* or externally supplied Cbl for propionate production (26, 34). Here, while investigating the cobamide metabolism of *A. muciniphila* strain Muc^T^, we uncovered a novel cobamide remodeling activity and identified and characterized an enzyme capable of initiating this process, CbiR. This discovery adds new complexity to the understanding of the roles of *A. muciniphila* in the gut. Not only does *A. muciniphila* degrade mucin to provide nutrients to the gut microbiota (26, 27), but it is also capable of altering cobamide structure, potentially changing the cobamide composition of its environment.

As a member of the AP endonuclease 2 superfamily, CbiR likely contains a (β/α)_8_ TIM barrel domain (63-77), unlike the structures predicted for the CbiZ and CobS protein families (80). Thus, not only does CbiR catalyze a unique reaction, but it is also distinct from the other cobamide remodeling enzymes in sequence and likely in structure. Intriguingly, while CbiR differs in sequence from B_12_-binding domains in cobamide-dependent enzymes, the substrate-binding domains of many cobamide-dependent enzymes are comprised of a (β/α)_8_ TIM barrel structure, with the C-terminal face interacting with the cobamide cofactor (81-91). Given that CbiR is predicted to have a similar fold, it is possible that cobamide binding in CbiR and in these cobamide-dependent enzymes shares common features. The yet to be discovered enzyme responsible for remodeling in algae may also be unique, as neither a *P. lutheri* transcriptome (92) nor the *C. reinhardtii* genome contains homologs of CbiR or CbiZ. It therefore appears that cobamide remodeling mechanisms have independently evolved multiple times. Together with the multiple pathways that exist for cobamide biosynthesis, transport, and precursor salvaging (31, 33, 35, 93-99), the addition of CbiR to the growing list of enzymes involved in cobamide metabolism highlights the importance of cobamide physiology in the evolution of bacteria.

CbiR has unexpectedly promiscuous activity, hydrolyzing cobamides irrespective of their lower ligand structure. This differs from *R. sphaeroides* CbiZ, which does not hydrolyze Cbl *in vitro* (59), and *V. cholerae* CobS, which remodels neither Cbl nor [Cre]Cba (48). In these cases, the cobamide remodeling pathway does not act on a cobamide(s) that can function in its organism’s metabolism. In contrast, *A. muciniphila* CbiR readily hydrolyzes pCbl, which functions as a cofactor for methionine synthesis and propionate metabolism in *A. muciniphila* and is the product of Cbi salvaging and cobamide remodeling in the bacterium itself. Thus, it is unclear how *A. muciniphila* prevents CbiR from continuing to hydrolyze pCbl after it is formed via cobamide remodeling. It is possible that pCbl is sequestered intracellularly by binding to MetH or other cobamide-dependent enzymes. Alternatively, CbiR activity could be coupled to cobamide uptake, as has been suggested for CbiZ (59, 100). Indeed, similar to some *cbiZ* homologs, 25% of *cbiR* homologs are located adjacent to genes for putative transport proteins, including in *A. muciniphila* strain Muc^T^ (Fig. S3). Remodeling in *D. mccartyi* strain 195 shows similar substrate promiscuity to *A. muciniphila* in the ability to act on numerous, structurally diverse cobamides (57), but the molecular basis of this promiscuity is unclear because its genome carries seven *cbiZ* homologs, none of which has been biochemically characterized. Aside from its activity on pCbl, the broad substrate range of CbiR may benefit *A. muciniphila* by enabling the bacterium to utilize a greater number of the cobamides present in its environment.

The discovery that *A. muciniphila* remodels cobamides leads us to reexamine its ecological roles in the gut. CbiR is found in all of the 75 recently sequenced *A. muciniphila* strains from the human and mouse gut, including the 26 strains that contain the *de novo* cobamide biosynthesis pathway (34, 101). Thus, like cobamide-dependent metabolism (34), cobamide remodeling appears to be nearly universal in *A. muciniphila*. Further, the role of *A. muciniphila* in the gut may be flexible, ranging from producing cobamides *de novo* to remodeling cobamides produced by other microbes, depending on which strains inhabit an individual. Notably, the end product of cobamide remodeling in *A. muciniphila*, pCbl, was the third most abundant corrinoid detected in the human gut in a study of human subjects residing at a single geographic location (42). Interestingly, that study also presented evidence that cobamide remodeling occurs in the human gut, as individuals supplemented with high levels of Cbl showed transiently increased levels of Cbi and the specific purinyl and phenolyl cobamides that were present in the gut prior to Cbl supplementation. It is possible that *A. muciniphila* is involved in this remodeling activity and contributes to the pool of pCbl in the gut. This, in turn, could modulate the growth or metabolism of other cobamide-requiring bacteria that rely on particular cobamides for their metabolic needs. CbiR may therefore not only expand access to the cobamides available to *A. muciniphila*, but also affect those accessible to other bacteria in the gut. Further, homologs of CbiR are found in at least 276 other microbial taxa and may function similarly in these microbes that inhabit diverse environments. The addition of CbiR to the cobamide remodeling enzymes that have been characterized to date – CbiZ, certain CobS homologs, and the enzyme(s) responsible for remodeling in algae – suggests that cobamide remodeling is more widespread than previously thought.

## Materials and Methods

### Media and growth conditions

*Akkermansia muciniphila* strain Muc^T^ (DSM 22959, ATCC BAA-835) was cultivated at 37°C in a vinyl anaerobic chamber (Coy Laboratory Products Inc) under an atmosphere of approximately 10% CO_2_, 3% H_2_, and 87% N_2_. A synthetic version of a basal mucin-based medium, in which mucin was replaced by soy peptone (16 g/L), L-threonine (4 g/L), glucose (2 g/L) and N-acetylglucosamine (2 g/L), was supplemented with 1% noble agar and used as a solid medium (21). This synthetic medium also contained L-methionine (125 mg/L) and omitted rumen fluid. M8 defined medium developed by Tramontano *et al*. (102) was used for liquid culturing. We found that the concentration of mucin in this medium (0.5%) was able to abrogate the requirement of methionine addition to the medium for *A. muciniphila* growth. Lowering the concentration to 0.25% resulted in methionine-deplete conditions for *A. muciniphila* and supplementation with methionine or cobamides restored robust growth. This mucin concentration was used for the MetH-dependent growth assays. However, batch to batch variations were seen with mucin such that media that supported robust growth while remaining methionine-deplete could not always be achieved. Cobamides were omitted from all growth media except when specified. For MetH-dependent growth assays, methionine was omitted from M8 medium.

*Escherichia coli* was cultured at 37°C with aeration in LB medium for cloning, protein expression, and assessing CbiR hydrolytic activity. Ethanolamine-based growth experiments used medium from Scarlett and Turner (38), with B_12_ omitted. Media were supplemented with antibiotics at the following concentrations when necessary: kanamycin, 25 mg/L (pETmini); ampicillin, 100 mg/L (pET-His_6_-MBP) and chloramphenicol, 20 mg/L (pLysS).

For all *in vivo* experiments involving corrinoids, culture media were supplemented with cyanylated cobamides or (CN)_2_Cbi, which are adenosylated following uptake into cells.

### Genetic and molecular cloning techniques

The entire *A. muciniphila cbiR* open reading frame, except the start codon, was cloned into a modified pET16b vector (103) with N-terminal His_6_ and MBP tags added for protein purification. For analysis of CbiR activity in *E. coli*, a minimized 3 kb derivative of pET28a (pETmini) containing the Kan^R^ marker, pBR322 origin, and *rop* gene was used. A constitutive promoter (BBa_J23100, iGEM) and RBS (BBa_B0034) were inserted into the vector for expression in *E. coli* MG1655-based strains (complete sequence: TTGACGGCTAGCTCAGTCCTAGGTACAGTGCTAGCGAATTCATACGACTCACTATAAAAGAGGAGAAA) and *A. muciniphila cbiR* was cloned downstream. Site-directed mutations were introduced into *cbiR* by PCR. All cloning was done by Gibson Assembly with *E. coli* XL1-Blue cells (104).

Construction of the *E. coli* MG1655 Δ*cobTSU* Δ*cobC* strain was accomplished using λ red-based recombination (105) and phage P1 transduction (106). An MG1655 Δ*cobTSU*::Kan^R^ operon deletion was constructed by λ red-based recombination. The Δ*cobC*::Kan^R^ allele was transduced into MG1655 via P1 transduction from *E. coli* strain JW0633-1, which was obtained from the Keio collection (107). Kan^R^ cassettes were removed by recombination of the flanking FRT sites as described (105).

### Chemical reagents

Porcine gastric mucin was purchased from MilliporeSigma (M1778). AdoCbl (coenzyme B_12_), MeCbl, CNCbl, and (CN)_2_Cbi were purchased from MilliporeSigma.

### Cobamide synthesis, adenosylation, and quantification

All other cyanylated cobamides used in the study were purified from bacterial cultures and cobamides were adenosylated and purified as previously described (50, 108). Cyanylated and adenosylated cobamides were quantified as previously described (50). MeCbl was quantified using an extinction coefficient of ε_519_ = 8.7 mM^-1^ cm^-1^ (109). AdoCbi-P and MeCbi-P were quantified using the dicyanylated corrinoid extinction coefficient ε_580_ = 10.1 mM^-1^ cm^-1^ following conversion to (CN)_2_Cbi-P by incubation with 10 mM KCN in the presence of light (110).

### *A. muciniphila* MetH-dependent growth assay

*A. muciniphila* was pre-cultured for 48 hours in M8 medium supplemented with 125 mg/L methionine. Cells were pelleted by centrifugation, washed twice with PBS, and diluted into 80 μL M8 medium to an OD_600_ of 0.02 in a 384-well plate (Nunc) with varying concentrations of cobamides and Cbi. The wells were sealed (ThermalSeal RTS(tm), Excel Scientific) and the plate was incubated at 37°C in a BioTek Epoch 2 microplate reader. OD_600_ was measured at regular intervals during growth.

### *E. coli* ethanolamine-dependent growth assay

*E. coli* was pre-cultured 16 h in ethanolamine medium supplemented with 0.02% ammonium chloride. Cells were pelleted by centrifugation, washed three times with 0.85% NaCl, and diluted to an OD_600_ of 0.025 in 200 μL ethanolamine medium with the specified cobamide additions in a 96-well plate (Corning). The wells were sealed (Breathe-Easy, Diversified Biotech) and OD_600_ was monitored at 37°C in a BioTek Synergy 2 microplate reader with shaking.

### Protein expression and purification

His_6_-MBP-CbiR was expressed in *E. coli* Rosetta(DE3) pLysS. Cells were grown to an OD_600_ of 0.4 at 37°C and expression was induced with 1 mM IPTG for 6 h at 30°C. Cells were lysed by sonication in 20 mM sodium phosphate, 300 mM NaCl, 10 mM imidazole, pH 7.4, with 0.5 mM PMSF, 1 μg/mL leupeptin, 1 μg/mL pepstatin, and 1 mg/mL lysozyme. The protein was purified from the clarified lysate using HisPur Ni-NTA resin (Thermo Scientific) and eluted with 250 mM imidazole. Purified protein was dialyzed into 25 mM Tris-HCl, pH 8.0, 300 mM NaCl, and 10% glycerol and stored at -80°C.

### His_6_-MBP-CbiR *in vitro* reactions

Due to light sensitivity of the compounds, all work involving adenosylated cobamides or MeCbl was performed in the dark or under red or dim white light. Unless specified, the *in vitro* reactions were performed anaerobically at 37°C in a vinyl anaerobic chamber with an atmosphere as described above. The components of the reactions were 50 mM Tris buffer, 0.3 μM purified His_6_-MBP-CbiR, 1 mM DTT, and variable concentrations of a cobamide. To prepare the Tris buffer, Tris base was dissolved and equilibrated within the anaerobic chamber. Prior to each experiment, the pH was adjusted with NaOH to account for acidification by the CO_2_ present in the atmosphere of the chamber. The pH was adjusted to 8.8-8.9 at room temperature (approximately 24°C), corresponding to a predicted pH of 8.45-8.55 at 37°C. Protein concentration was determined by absorbance at 280 nm (A_280_).

A BioTek Epoch 2 microplate reader and half-area UV-Star® 96-well microplates (Greiner Bio-One) were used for assays monitoring the reaction by absorbance. For these assays, separate 2X solutions of AdoCbl and His_6_-MBP-CbiR were prepared in 50 mM Tris buffer and 1 mM DTT. A frozen aliquot of His_6_-MBP-CbiR was thawed inside the anaerobic chamber prior to dilution. The 2X AdoCbl solution was pre-incubated at 37°C for 60 min, while the 2X CbiR solution was pre-incubated at 37°C for 20 min. 60 μL each of 2X AdoCbl and 2X His_6_-MBP-CbiR were then mixed in a 96-well plate, with 100 μL transferred to a new well prior to measurements in the plate reader. Assays with MeCbl were prepared similarly.

Absorbances over time at 534 and 527 nm were used to monitor the conversion of AdoCbl to AdoCbi-P and MeCbl to MeCbi-P, respectively. To enable conversion of A_534_ into moles of AdoCbl and A_527_ into moles of MeCbl, the extinction coefficients of AdoCbl, MeCbl, and purified AdoCbi-P and MeCbi-P were determined in 50 mM Tris buffer, pH 8.8, 1 mM DTT: ε_534_ (AdoCbl) = 7.8 mM^-1^ cm^-1^, ε_527_ (MeCbl) = 8.0 mM^-1^ cm^-1^, ε_534_ (AdoCbi-P) = 1.3 mM^-1^ cm^-1^, ε_527_ (MeCbi-P) = 2.7 mM^-1^ cm^-1^.

For reactions with adenosylated cobamides with different lower ligands monitored by HPLC, cobamides were mixed at 60 μM with 50 mM Tris buffer and 1 mM DTT and equilibrated to 37°C. His_6_-MBP-CbiR was equilibrated to 37°C at 0.6 μM in 50 mM Tris buffer and 1 mM DTT. Each cobamide and His_6_-MBP-CbiR were mixed in equal volume and incubated at 37°C. At three different timepoints, 100 μL of the reaction mix was removed and mixed with 5 μL of 600 mM EDTA to quench the reaction. The protein was removed from samples using Nanosep 10K centrifugal devices (Pall) prior to injection onto the HPLC. AdoCbi-P levels in the samples were quantified by HPLC by comparing to a standard curve generated with known quantities of purified AdoCbi-P. For reactions involving incubations of 4-18 h, initial equilibration at 37°C was not performed.

### Corrinoid extraction

Cbi salvaging and cobamide remodeling were assessed in *A. muciniphila* by cultivating in M8 medium supplemented with 10 nM Cbi or cobamide, respectively, for 72 h. CbiR cobamide hydrolytic activity with different cobamides in *E. coli* was monitored using the MG1655 Δ*cobTSU* Δ*cobC* strain cultivated in LB medium supplemented with 10 nM cobamide for 16 h. Cobamide remodeling in *E. coli* MG1655 was assessed by cultivating in ethanolamine medium supplemented with 100 nM [Cre]Cba and 1 μM DMB for 94 h. CbiR mutants were analyzed in *E. coli* MG1655 Δ*cobTSU* Δ*cobC* by culturing in LB medium supplemented with 75 nM Cbl for 20 h.

Cyanation of corrinoids extracted from cells for Figures 2A, 2C, S1, and S2 was performed as previously described (57), with 5,000 corrinoid molar equivalents of KCN added. For extractions of adenosylated corrinoids (Figures 5A, 6C, S5, and S6), cell lysis was performed similarly, with KCN omitted; following removal of cellular debris by centrifugation, deionized water was added to the supernatant to decrease the methanol concentration to 10%. Solid phase extraction of cyanylated and adenosylated corrinoids with Sep-Pak C18 cartridges (Waters) was performed as described (37). Samples were dried, resuspended in 200 μL deionized water (pH 7), and filtered with 0.45 μm filters (Millex-HV, Millipore) or Nanosep 10K centrifugal devices prior to analysis by HPLC. For extractions involving adenosylated cobamides, all steps were performed in the dark or under red or dim white light.

### HPLC and MS analysis

Corrinoids were analyzed on an Agilent 1200 series HPLC equipped with a diode array detector. For experiments in Figures 2A, 2C, 3B, 4A, and 4B, an Agilent Zorbax SB-Aq column (5 μm, 4.6 ⨯ 150 mm) was used as previously described (method 2, (58)). For experiments in Figures 5A, 6C, and S6A, an Agilent Zorbax Eclipse Plus C18 column (5 μm, 9.4 × 250 mm) was used with the following method: Solvent A, 0.1% formic acid in deionized water; Solvent B, 0.1% formic acid in methanol; 2 mL/min at 30°C; 18% solvent B for 2.5 min followed by a linear gradient of 18 to 60% solvent B over 28.5 min.

An Agilent 1260 series fraction collector was used for HPLC purification of corrinoids and CbiR reaction products. The purification of CN-pCbl from *A. muciniphila* was performed using the Zorbax SB-Aq column with the method described above. Purification of AdoCbi-P and α-ribazole from the hydrolysis of AdoCbl by His_6_-MBP-CbiR was performed in two steps. AdoCbi-P and α-ribazole were first separated and collected on a Zorbax Eclipse XDB-C18 column (5 μm, 4.6 × 150 mm) using the following method: Solvent A, 10 mM ammonium acetate pH 6.5; Solvent B, 100% methanol; 1 mL/min at 30°C; 0% B for 2 min followed by a linear gradient of 0 to 15% solvent B over 1.5 min, 15 to 35% over 6.5 min, 35 to 70% over 2 min, and 70 to 100% over 2 min. Each compound was subsequently run and collected on the Zorbax SB-Aq column with the method described above. The purification of MeCbi-P from the *in vitro* hydrolysis of MeCbl was performed using the Zorbax SB-Aq column with the method above. The purification of adenosylated hydrolysis products of CbiR from *E. coli* was performed with two rounds of collection; AdoCbi-P and AdoCbi were first separated and collected on the Zorbax Eclipse Plus C18 column with the method above, and then each compound was run and collected on the Zorbax SB-Aq column using the method described. AdoCbl remodeled from [Cre]Cba in *E. coli* was purified using the Zorbax Eclipse Plus C18 column with the method described above. Collected compounds were de-salted with Sep-Pak C18 cartridges.

MS analysis was performed on a Bruker Linear Iontrap Quadrupole coupled to a Fourier Transform Ion Cyclotron (LTQ-FT) mass spectrometer at the QB3/Chemistry MS Facility (UC Berkeley). Prior to MS analysis, the purified adenosylated and methylated corrinoids were exposed to light to remove the adenosyl and methyl upper ligands, respectively.

### Phylogenetic analysis

282 homologs of *A. muciniphila* strain Muc^T^ CbiR were identified by BLAST with Expect values lower than 10^−3^ (Table S1; sequences from other strains of *A. muciniphila* are excluded). A subset of 203 sequences with Expect values lower than 10^−14^ and whose encoding genes were located adjacent to and in the same orientation as a cobamide biosynthesis gene(s) were chosen for phylogenetic analysis with the AP endonuclease 2 superfamily (pfam 01261) (Table S1). Sequences were clustered at 0.95 using CD-HIT to reduce the CbiR homolog sequence set by removing subspecies sequence diversity (111). This final set of 178 sequences and *A. muciniphila* CbiR were used to infer a phylogenetic tree with experimentally characterized members of the AP endonuclease 2 superfamily (Table S2). The sequences were aligned with MAFFT (112) and positions with 95% or greater gaps were removed by trimAl (113). A maximum likelihood tree presented in Figure 6A was inferred from this alignment using IQ-TREE v1.6.12 (114) with 1,500 ultrafast bootstraps and visualized in iTOL (115).

## Acknowledgements

We thank Amanda Shelton, Amrita Hazra, Kristopher Kennedy, Per Malkus, Raphael Valdivia, and Sebastian Gude for helpful discussions and Kristopher Kennedy and Sebastian Gude for critical reading of the manuscript. We are grateful to Kristopher Kennedy and Sebastian Gude for construction of the pETmini plasmid and *E. coli* MG1655 Δ*cobTSU* Δ*cobC* strain, Nina Kirmiz and Gilberto Flores for reagents, and Rita Nichiporuk at the UC Berkeley QB3 Mass Spectrometry Facility for analysis of MS samples. This work was funded by National Institutes of Health Grant R01GM114535.

---

**Figure S1.**
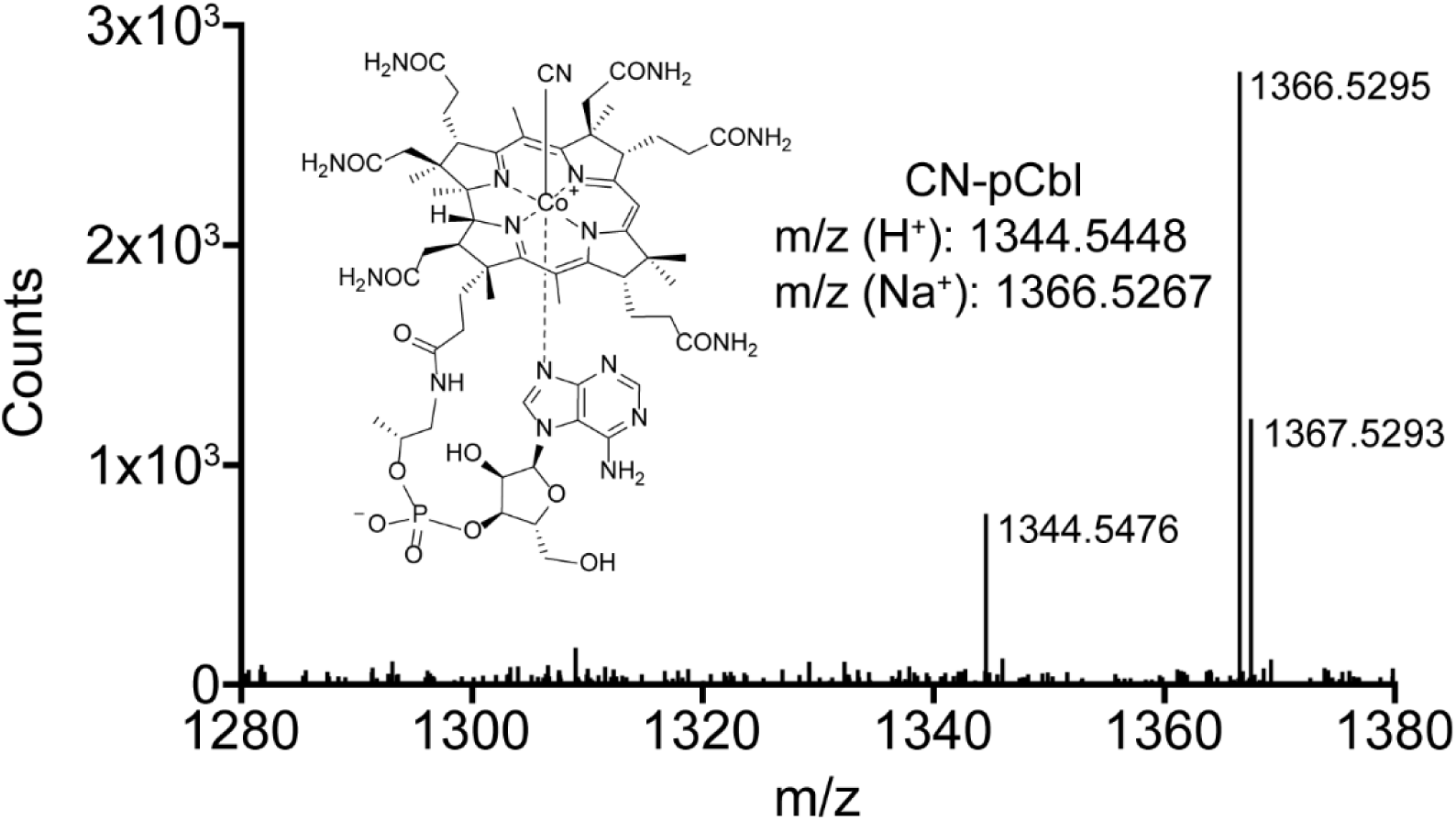
The corrinoid produced by *A. muciniphila* when grown with Cbi was purified by HPLC and analyzed by MS. The structure and predicted m/z for CN-pCbl are shown for comparison.

**Figure S2.**
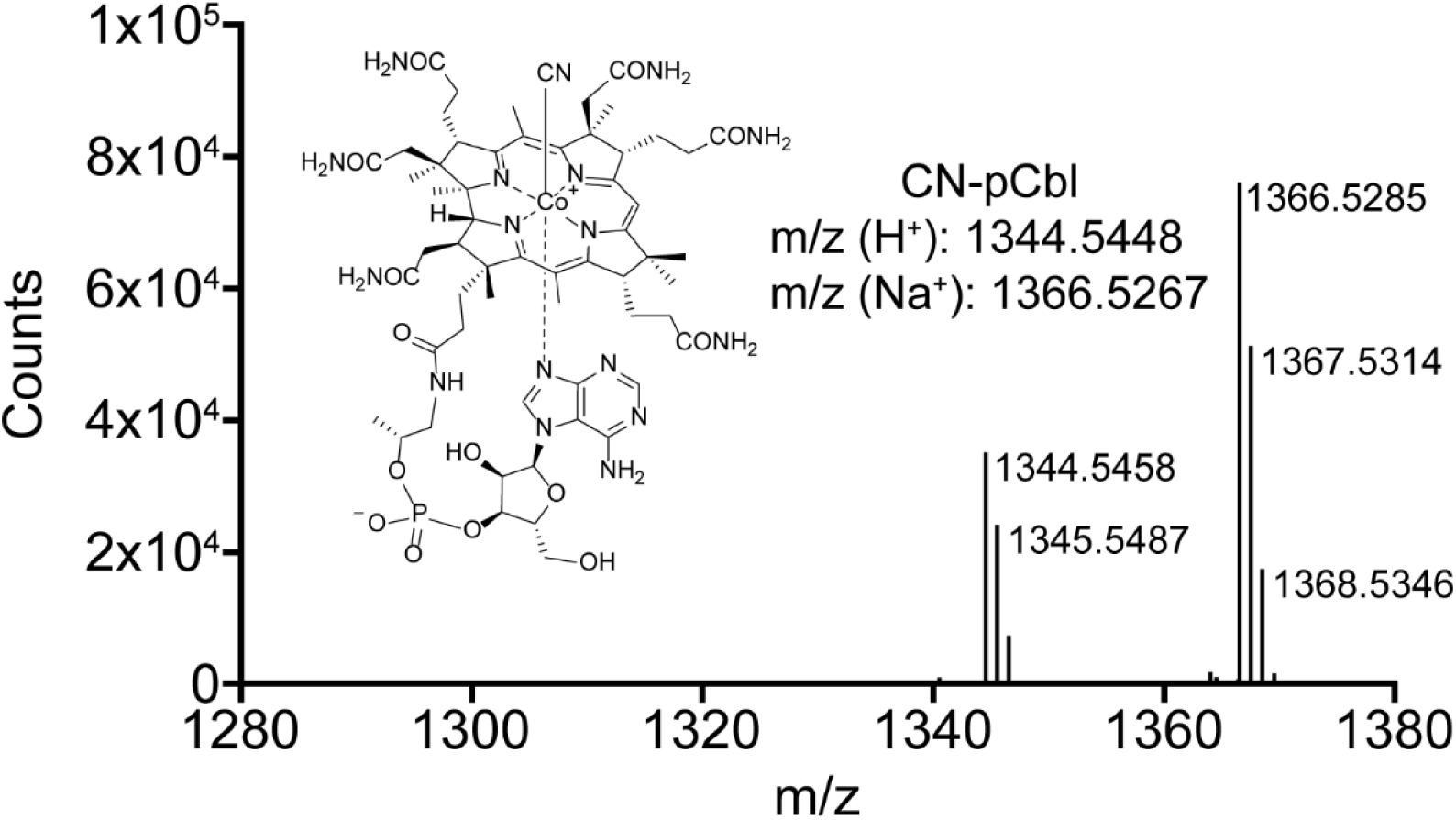
The corrinoid produced by *A. muciniphila* when grown with Cbl was purified by HPLC and analyzed by MS. The structure and predicted m/z for CN-pCbl are shown for comparison.

**Figure S3.**
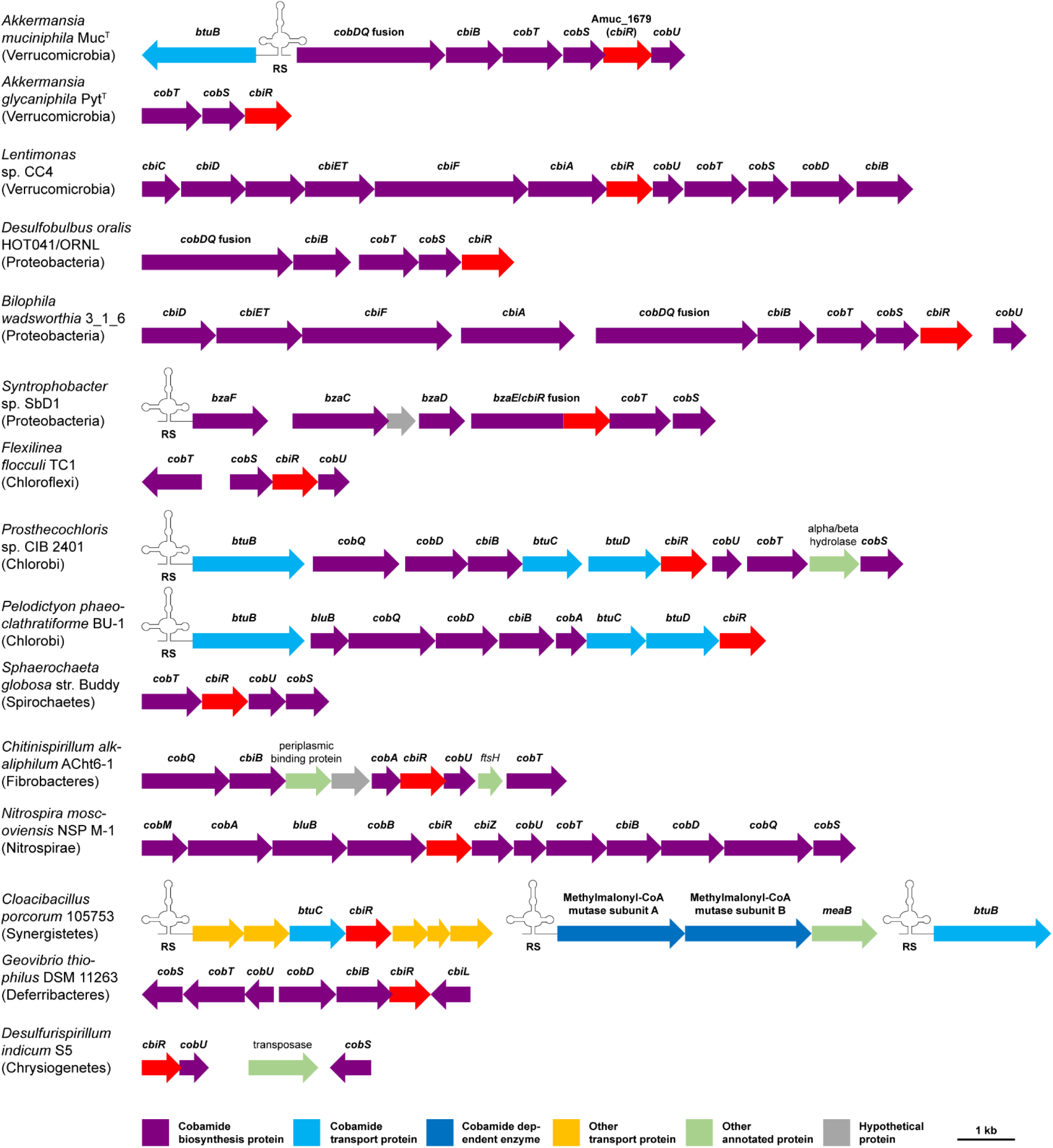
Expanded list of homologs of Amuc_1679 (*cbiR*, red arrows). Species and strain names are given, with phylum names in parentheses. RS denotes a predicted cobalamin riboswitch. The lengths and positions of the ORFs are drawn to scale.

**Figure S4.**
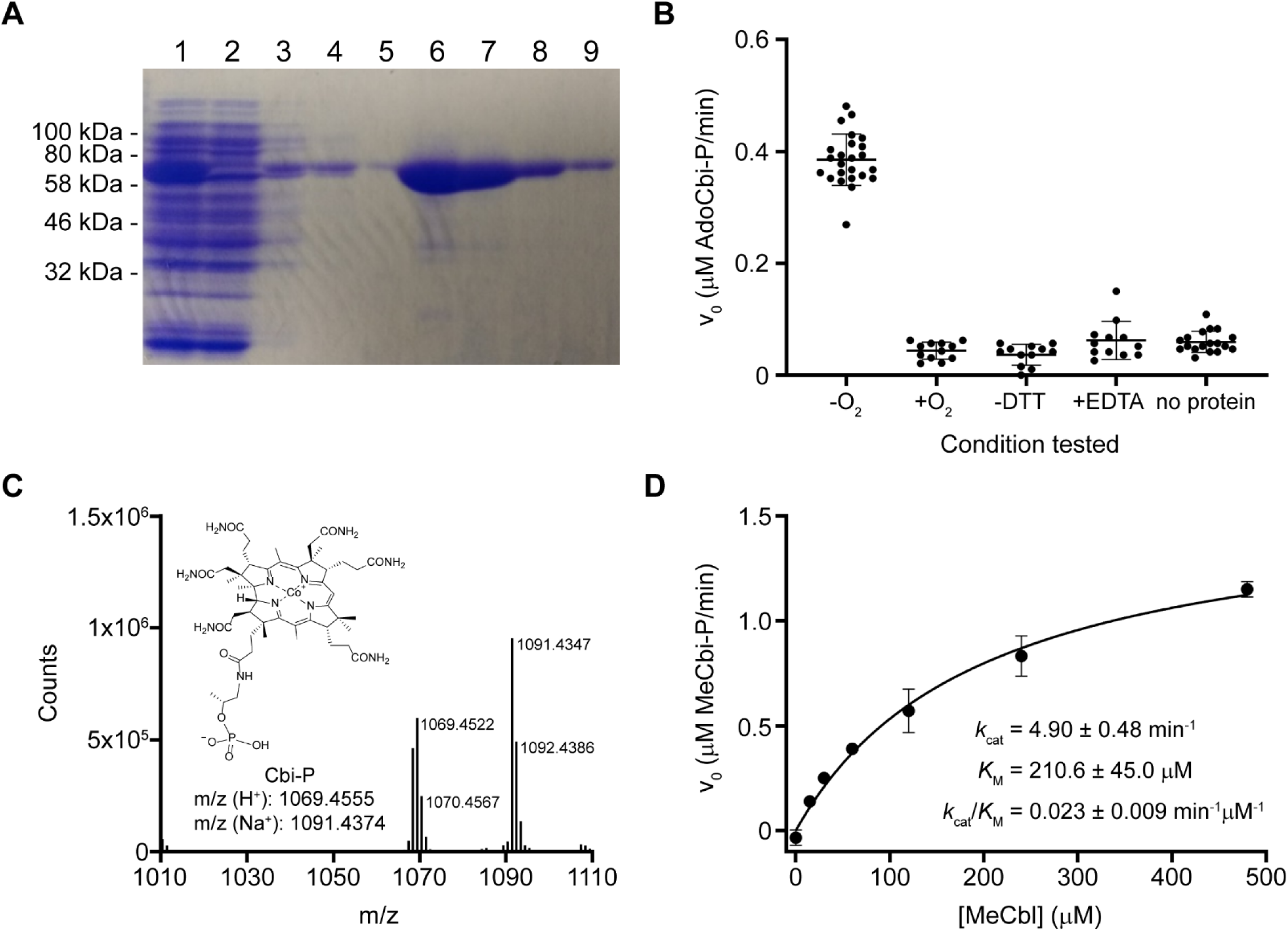
Biochemical characterization of His_6_-MBP-CbiR. A. SDS-PAGE gel of purification of His_6_-MBP-CbiR with Ni-NTA resin. Lane 1: Cell lysate, Lane 2: Flowthrough, Lanes 3-5: Wash fractions, Lanes 6-9: Elution fractions. His_6_-MBP-CbiR has a predicted molecular weight of 77 kDa. *B. In vitro* characterization of His_6_-MBP-CbiR. The reaction rates were determined for 0.3 µM His_6_-MBP-CbiR incubated with 30 µM AdoCbl. The standard reaction mixture used throughout the manuscript (labeled as -O_2_) contained 50 mM Tris, pH 8.45-8.55 and 1 mM DTT and was performed in an anaerobic chamber. Activity was also measured in ambient O_2_ (+O_2_), with DTT omitted (-DTT), with 3 µM EDTA added (+EDTA), and in the absence of His_6_-MBP-CbiR (no protein). The reaction was monitored by measuring A_534_ over time. Lines and error bars show the mean and standard deviation, respectively. C. The corrinoid product of His_6_-MBP-CbiR incubated with MeCbl was purified by HPLC, exposed to light to remove the methyl upper ligand, and analyzed by MS. The structure and predicted m/z for Cbi-P are shown for comparison. D. Michaelis-Menten kinetic analysis of His_6_-MBP-CbiR with MeCbl. His_6_-MBP-CbiR was tested at 0.3 µM and the reaction was monitored by measuring the decrease in absorbance at 527 nm (A_527_). The experiment was performed twice with similar results. Data from a representative experiment with three technical replicates are shown. Kinetic constants were calculated using data from all six replicates.

**Figure S5.**
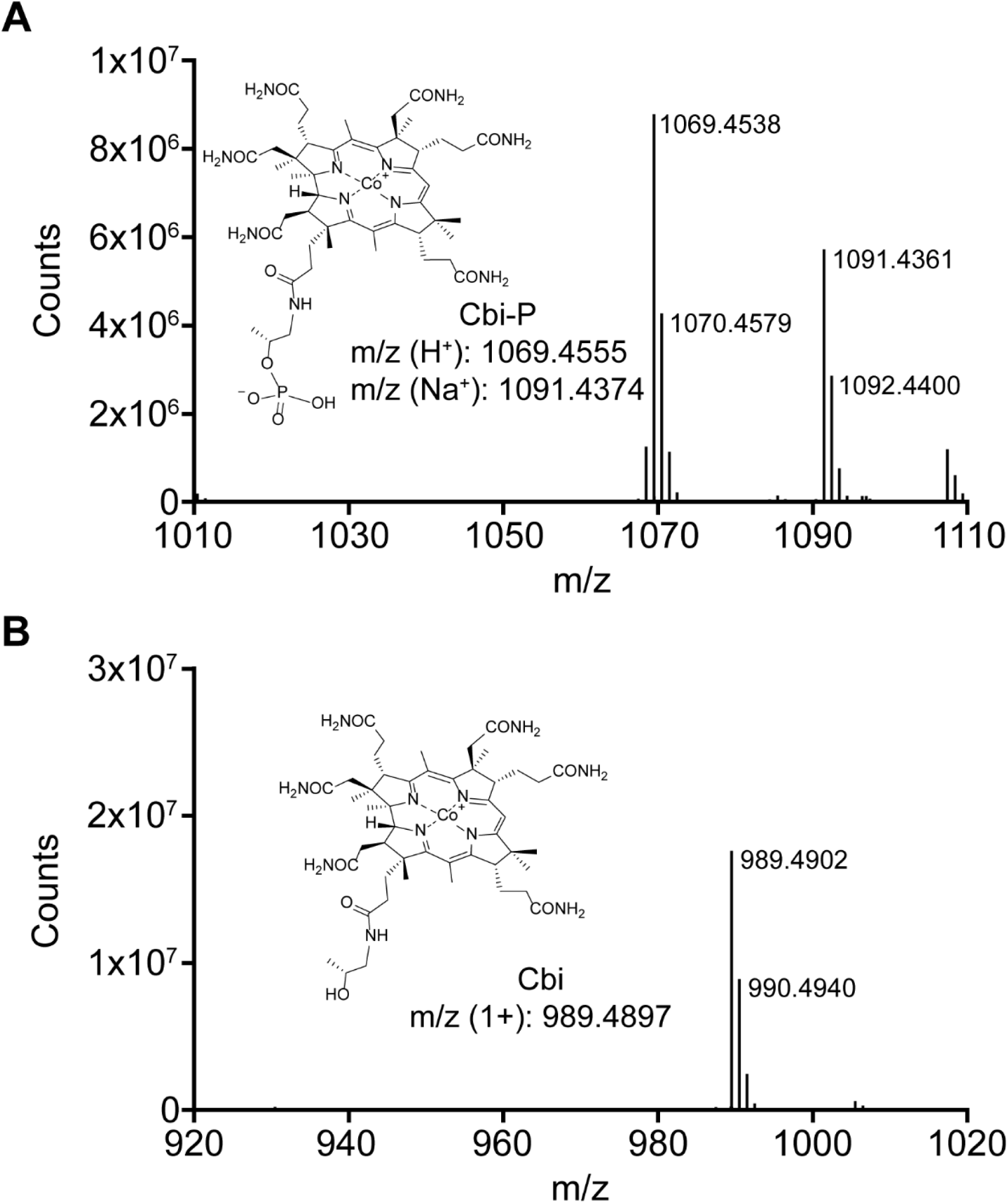
MS analysis of CbiR hydrolysis products extracted from *E. coli*. MS analysis of the corrinoid products labeled with asterisks in Fig. 5A are shown for the peaks at A. 16 min and B. 18 min. The corrinoids were purified by HPLC and exposed to light to remove the adenosyl upper ligand prior to analysis by MS. The structures and predicted m/z for Cbi-P and Cbi are shown for comparison.

**Figure S6.**
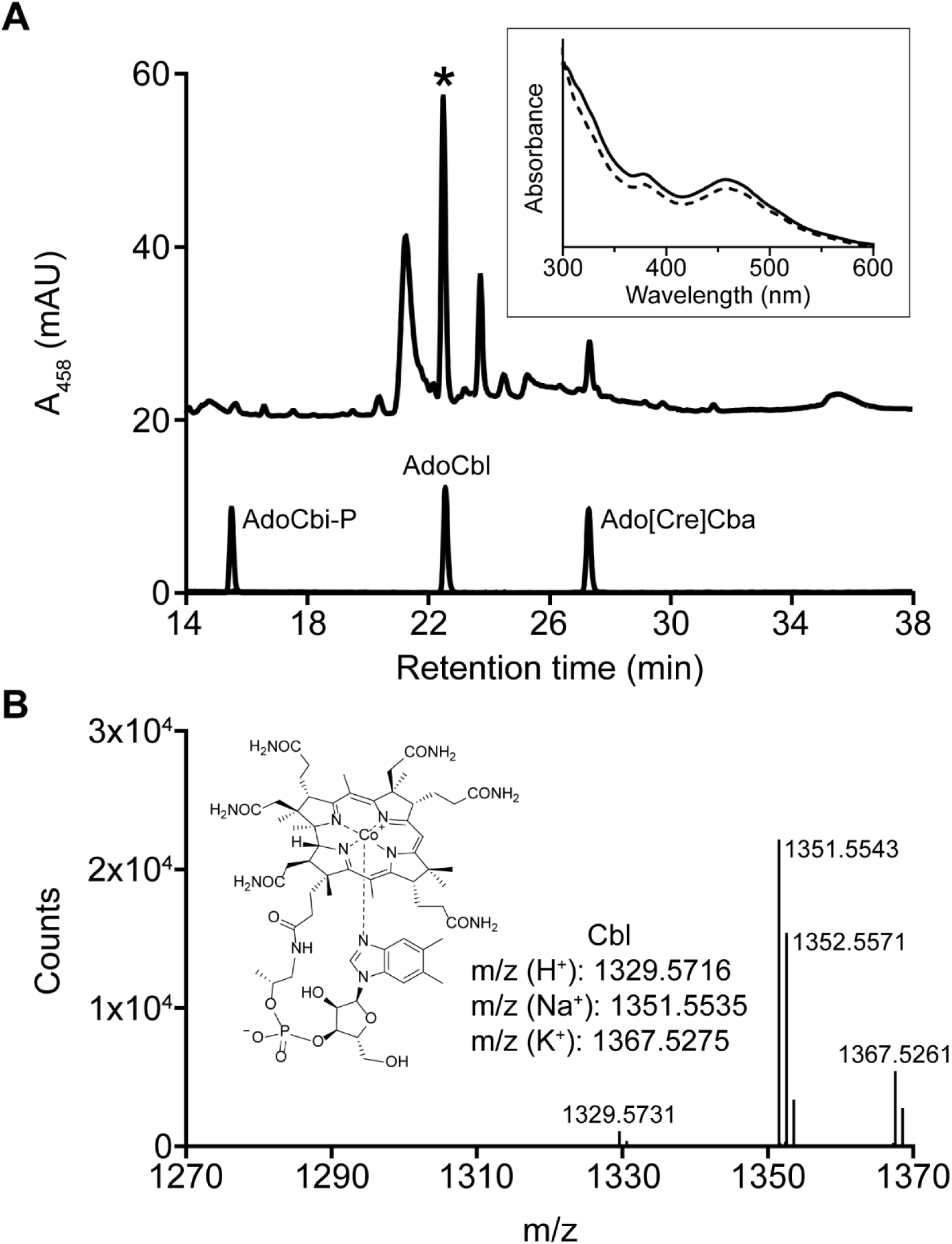
Cobamide remodeling of [Cre]Cba to Cbl in *E. coli* expressing *cbiR*. A. Remodeling of [Cre]Cba to Cbl in *E. coli*. Wild type strain MG1655 containing pETmini-*cbiR* was grown in 1L minimal medium with ethanolamine supplemented with 100 nM [Cre]Cba and 1 µM DMB. Corrinoids were extracted and analyzed by HPLC. AdoCbl and Ado[Cre]Cba standards and purified AdoCbi-P are shown at the bottom. Comparison of the UV-Vis spectra of AdoCbl (dashed line) and the starred peak (solid line) are shown in the inset; spectra were normalized to each other at the local maxima at 458 nm to aid comparison. The spectrum of AdoCbl differs from that in Fig. 3E due to the acidic HPLC conditions. B. MS analysis of the peak labeled with an asterisk in panel A. The corrinoid was purified by HPLC and exposed to light to remove the adenosyl upper ligand prior to analysis by MS. The structure and predicted m/z for Cbl are shown for comparison.

**Table S1.** Homologs of A. muciniphila CbiR

| Organism | Phylum | Isolation Source | Accession number | E value | % Identity to *A. muciniphila* MucT CbiR | Adjacent to and in same orientation as cobamide biosynthesis gene(s)? |
| --- | --- | --- | --- | --- | --- | --- |
| Akkermansia muciniphila strain MucT | Verrucomicrobia | human feces | WP_081429195.1 |  | (100) | Y |
| Akkermansia glycaniphila strain Pyt | Verrucomicrobia | reticulated python feces | OCA02218.1 | 2.E-115 | 62 | Y |
| Akkermansia sp. strain BIOML-A47 | Verrucomicrobia | human feces | KAA3306648.1 | 2.E-174 | 84 | Y |
| Akkermansia sp. strain BIOML-A60 | Verrucomicrobia | human feces | KAA3165711.1 | 1.E-175 | 85 | Y |
| Akkermansia sp. strain KLE1797 | Verrucomicrobia | human feces | KXT49684.1 | 7.E-173 | 87 | Y |
| Aminomonas paucivorans strain DSM 12260 | Synergistetes | anaerobic lagoon of a dairy wastewater treatment plant (Columbia) | EFQ22869.1 | 9.E-15 | 28 | N |
| anaerobic bacterium MO-CFX2 | Chloroflexi | marine sediment (Shimokita Peninsula, Japan) | WP_162909697.1 | 5.E-27 | 31 | Y |
| Anaerolineaceae bacterium isolate AS05jafATM_106 | Chloroflexi | anaerobic digestion of organic wastes under variable temperature conditions and feedstocks | HHV05068.1 | 5.E-35 | 35 | Y |
| Anaerolineaceae bacterium oral taxon 439 strain W11661 | Chloroflexi | human subgingival dental plaque | AOH42763.1 | 7.E-35 | 35 | Y |
| Anaerolineae bacterium isolate AS06rmzACSIP_331 | Chloroflexi | anaerobic digestion of organic wastes under variable temperature conditions and feedstocks | NLF10836.1 | 4.E-24 | 30 | Y |
| Anaerolineae bacterium isolate HyVt-475 | Chloroflexi | rock sample from hydrothermal vent (Lau Basin, Tonga) | HFD38495.1 | 6.E-23 | 30 | Y |
| Anaerolineae bacterium isolate Nak57 | Chloroflexi | sulfidic hot spring (Azumino, Japan) | PWH15486.1 | 3.E-37 | 34 | Y |
| Anaerolineae bacterium isolate SpSt-822 | Chloroflexi | mud, sediment (Yellowstone National Park, USA) | HGX29340.1 | 2.E-25 | 30 | Y |
| Anoxybacter fermentans strain DY22613 | Firmicutes | deep-sea hydrothermal vent | AZR72222.1 | 1.E-04 | 25 | N |
| Ardenticatena maritima strain 110S | Chloroflexi | hydrothermal field (Yamagawa, Japan) | GAP62401.1 | 3.E-16 | 30 | Y |
| Bellilinea caldifistulae isolate SpSt-556 | Chloroflexi | hot spring sediment (British Columbia, Canada) | HGS87922.1 | 2.E-30 | 33 | Y |
| Bilophila sp. strain 4_1_30 | Proteobacteria | human feces | EGW44733.1 | 5.E-66 | 41 | Y |
| Bilophila wadsworthia strain 3_1_6 | Proteobacteria | human feces | EFV46162.1 | 2.E-64 | 40 | Y |
| Burkholderiales bacterium isolate AS22ysBPME_216 | Proteobacteria | anaerobic digestion of organic wastes under variable temperature conditions and feedstocks | NMA29311.1 | 1.E-14 | 28 | Y |
| Caldilinea aerophila isolate SpSt-289 | Chloroflexi | hot spring sediment (British Columbia, Canada) | HDX32185.1 | 2.E-24 | 28 | Y |
| Caldilinea aerophila strain DSM 14535 | Chloroflexi | sulfur-turf sample in a hot spring (Takayama, Japan) | BAL99387.1 | 4.E-24 | 28 | N |
| Calditerrivibrio nitroreducens strain DSM 19672 | Deferribacteres | terrestrial hot spring (Lake Nojiri, Japan) | ADR19788.1 | 3.E-05 | 26 | Y |
| candidate division KSB1 bacterium isolate B7_G2 | Candidate division KSB1 | deep-sea hydrothermal vent sediment (Guaymas Basin, Mexico) | RKY75023.1 | 6.E-16 | 26 | Y |
| candidate division KSB1 bacterium isolate B39_G15 | Candidate division KSB1 | deep-sea hydrothermal vent sediment (Guaymas Basin, Mexico) | RKY84749.1 | 7.E-16 | 26 | Y |
| candidate division KSB1 bacterium isolate CS2-K093 | Candidate division KSB1 | pelagic sediment (Gulf of Kutch, India) | NIV91984.1 | 2.E-25 | 33 | Y |
| Candidatus Abyssobacterium bacterium isolate SURF_5 | Candidatus Abyssobacterium | deeply circulating subsurface aquifer fluids (Lead, SD, USA) | RJP21339.1 | 7.E-07 | 27 | N |
| Candidatus Acidulodesulfobacterium acidiphilum isolate AP4 | Proteobacteria | acid mine drainage (Guangzhou, China) | RZV40568.1 | 3.E-05 | 31 | N |
| Candidatus Acidulodesulfobacterium ferriphilum isolate AP3 | Proteobacteria | acid mine drainage (Guangzhou, China) | RZD14164.1 | 2.E-04 | 27 | N |
| Candidatus Fermentibacter daniensis isolate Ran_1 | Candidatus Fermentibacteria | enrichment reactor seeded with activated sludge (Randers, Denmark) | KZD16791.1 | 8.E-25 | 31 | Y |
| Candidatus Handelsmanbacteria bacterium isolate RIFCSPLOWO2_12_FULL_64_10 | Candidatus Handelsmanbacteria | Rifle well FP-101 (Rifle, CO, USA) | OGG49119.1 | 3.E-28 | 34 | Y |
| Candidatus Latescibacteria bacterium isolate UBA9260 | Candidatus Latescibacteria | marine | HAA78539.1 | 2.E-28 | 32 | Y |
| Candidatus Methyloiridis limnetica strain Zug | candidate division NC10 | freshwater (Lake Zug, Switzerland) | PTL35634.1 | 4.E-21 | 32 | Y |
| Candidatus Methyloiridis oxyfera isolate SB1 | candidate division NC10 | biomass on anode of bioreactor after 21 days (Ooijpolder, Netherlands) | KAB2960242.1 | 8.E-18 | 30 | Y |
| Candidatus Nitrosotenuis sp. isolate GW928_bin.18 | Thaumarchaeota | groundwater (Oak Ridge, TN, USA) | TBR21516.1 | 4.E-16 | 28 | Y |
| Candidatus Raymondobacteria bacterium isolate RifOxyA12_full_50_37 | Candidatus Raymondobacteria | Rifle well CD01 (Rifle, CO, USA) | OGJ89256.1 | 8.E-16 | 29 | Y |
| Candidatus Thermofonsia Clade 1 bacterium isolate CP2_2F | Chloroflexi | alkaline, sulfidic hot spring Cone Pool 2 (Azumino, Japan) | PJF30363.1 | 4.E-42 | 36 | Y |
| Candidatus Woesebacteria bacterium isolate B4_G12 | Candidatus Woesebacteria | deep-sea hydrothermal vent sediments (Guaymas Basin, Mexico) | RLC29799.1 | 1.E-22 | 30 | Y |
| Chitinispirillum alkaliphilum strain AChT6-1 | Fibrobacteres | hypersaline alkaline lake (Wadi al Natrun, Egypt) | KMQ49640.1 | 1.E-19 | 29 | Y |
| Chlorobaculum limnaeum strain DSM1677 | Chlorobi | Lake Kinnevet, Israel | AOS83552.1 | 3.E-22 | 32 | Y |
| Chlorobaculum parvum strain NCIB 8327 DSM 263 | Chlorobi | freshwater | ACF11581.1 | 3.E-19 | 31 | Y |
| Chlorobaculum sp. strain 24CR | Chlorobi | water (Carmel River, CA, USA) | RXK88351.1 | 3.E-21 | 31 | Y |
| Chlorobaculum tepidum strain TLS | Chlorobi | high-sulfide hot spring (Rotorua, New Zealand) | AAM72179.1 | 1.E-17 | 29 | Y |
| Chlorobaculum thiosulfatophilum strain DSM 249 | Chlorobi | tassajara hot spring (CA, USA) | TNJ40258.1 | 2.E-22 | 31 | Y |
| Chlorobiaceae bacterium isolate June25_Bin_2 | Chlorobi | freshwater lake (Trout Bog Lake, WI, USA) | NMW21650.1 | 4.E-17 | 27 | Y |
| Chlorobiaceae bacterium isolate Oct13_Bin_1 | Chlorobi | freshwater lake (Trout Bog Lake, WI, USA) | NMW18963.1 | 6.E-16 | 28 | Y |
| Chlorobium chlorochromatii strain CaD3 | Chlorobi | eutrophic lake (Brandenburg, Germany) | ABB28338.1 | 6.E-16 | 30 | N |
| Chlorobium ferrooxidans strain DSM 13031 | Chlorobi | ditch sediment (Konstanz, Germany) | EAT58859.1 | 6.E-17 | 27 | Y |
| Chlorobium limicola strain DSM 245 | Chlorobi | hot spring (CA, USA) | ACD90141.1 | 5.E-17 | 29 | Y |
| Chlorobium limicola strain Frasassi | Chlorobi | sediment-water interface in an artificial aquarium in the Frasassi cave system (Italy) | KUL32430.1 | 3.E-16 | 29 | N |
| Chlorobium phaeobacteroides strain BS1 | Chlorobi | chemocline of the Black Sea | ACE04425.1 | 3.E-17 | 29 | N |
| Chlorobium phaeobacteroides strain DSM 266 | Chlorobi | Lake Blankvann (Norway) | ABL65163.1 | 3.E-24 | 33 | Y |
| Chlorobium phaeovibrioides strain BrKhr17 | Chlorobi | Lake Bolshye Khruslomeny, water chemocline zone | RTY40038.1 | 7.E-18 | 31 | Y |
| Chlorobium phaeovibrioides strain DSM 265 | Chlorobi | saline intertidal flat | ABP36835.1 | 7.E-20 | 31 | Y |
| Chlorobium phaeovibrioides strain PhvTcv-s14 | Chlorobi | water chemocline zone (Russia) | QEQ57063.1 | 6.E-18 | 31 | Y |
| Chlorobium sp. isolate L227-2013-22 | Chlorobi | freshwater lake (Kenora, Ontario, Canada) | TLU55211.1 | 2.E-21 | 31 | Y |
| Chlorobium sp. isolate L227-2013-55 | Chlorobi | freshwater lake (Kenora, Ontario, Canada) | TLU57894.1 | 8.E-19 | 29 | N |
| Chlorobium sp. isolate L227-2013-56 | Chlorobi | freshwater lake (Kenora, Ontario, Canada) | TLU51735.1 | 9.E-26 | 30 | Y |
| Chlorobium sp. isolate L227-S-6D | Chlorobi | freshwater lake (Kenora, Ontario, Canada) | TLU81745.1 | 6.E-17 | 29 | N |
| Chlorobium sp. isolate L304-S-6D | Chlorobi | freshwater lake (Kenora, Ontario, Canada) | TLU83342.1 | 2.E-21 | 31 | Y |
| Chlorobium sp. isolate L442-64 | Chlorobi | freshwater lake (Kenora, Ontario, Canada) | TLU54122.1 | 4.E-18 | 29 | N |
| Chlorobium sp. isolate UBA8843 | Chlorobi | synthetic microbial community | HCD35528.1 | 9.E-20 | 31 | Y |
| Chlorobium sp. strain BLA1 | Chlorobi | Brownie Lake (Minneapolis, MN, USA) | NHQ60379.1 | 2.E-17 | 28 | Y |
| Chlorobium sp. strain KB01 | Chlorobi | meromictic lake (Kabuno Bay, Congo) | WP_076791818.1 | 4.E-13 | 27 | Y |
| Chlorobium sp. strain N1 | Chlorobi | sediment (Norsminde Fjord, Denmark) | TCD47456.1 | 8.E-21 | 30 | Y |
| Chloroflexi bacterium isolate AS06rmzACSIP_450 | Chloroflexi | anaerobic digestion of organic wastes under variable temperature conditions and feedstocks | NLW71729.1 | 7.E-33 | 32 | Y |
| Chloroflexi bacterium isolate AS15tH2ME_173 | Chloroflexi | anaerobic digestion of organic wastes under variable temperature conditions and feedstocks | NLA79515.1 | 6.E-35 | 34 | Y |
| Chloroflexi bacterium isolate AS26fmACSIPLY_11 | Chloroflexi | anaerobic digestion of organic wastes under variable temperature conditions and feedstocks | HHY58046.1 | 2.E-22 | 30 | Y |
| Chloroflexi bacterium isolate AS27yjCOA_1 | Chloroflexi | anaerobic digestion of organic wastes under variable temperature conditions and feedstocks | NMB61392.1 | 1.E-36 | 31 | Y |
| Chloroflexi bacterium isolate AS27yjCOA_123 | Chloroflexi | anaerobic digestion of organic wastes under variable temperature conditions and feedstocks | NMB86957.1 | 4.E-33 | 33 | Y |
| Chloroflexi bacterium isolate AS27yjCOA_124 | Chloroflexi | anaerobic digestion of organic wastes under variable temperature conditions and feedstocks | NMB61392.1 | 1.E-36 | 31 | Y |
| Chloroflexi bacterium isolate AS27yjCOA_4 | Chloroflexi | anaerobic digestion of organic wastes under variable temperature conditions and feedstocks | NMC52453.1 | 1.E-23 | 33 | Y |
| Chloroflexi bacterium isolate AS27yjCOA_56 | Chloroflexi | anaerobic digestion of organic wastes under variable temperature conditions and feedstocks | NMC45596.1 | 4.E-35 | 32 | Y |
| Chloroflexi bacterium isolate B46_G1 | Chloroflexi | deep-sea hydrothermal vent sediment (Guaymas Basin, Mexico) | RLC71331.1 | 5.E-39 | 35 | Y |
| Chloroflexi bacterium isolate CFX1 | Chloroflexi | marine ANAMMOX bioreactor | KAA3663928.1 | 4.E-35 | 34 | N |
| Chloroflexi bacterium isolate CR04 | Chloroflexi | marine sediment (Pacific Ocean) | TDI86467.1 | 2.E-33 | 33 | Y |
| Chloroflexi bacterium isolate HGW-Chloroflexi-1 | Chloroflexi | groundwater (Horonobe, Japan) | PKO23290.1 | 1.E-33 | 34 | Y |
| Chloroflexi bacterium isolate HGW-Chloroflexi-10 | Chloroflexi | groundwater (Horonobe, Japan) | PKO13937.1 | 1.E-35 | 34 | Y |
| Chloroflexi bacterium isolate M_MetaBat.114 | Chloroflexi | deepsea hydrothermal sulfide chimney (Pacific Ocean) | NOZ27705.1 | 8.E-31 | 30 | Y |
| Chloroflexi bacterium isolate metabat2.725 | Chloroflexi | Prairie Pothole Region wetland sediment (Cottonwood Lake, ND, USA) | RPH61221.1 | 1.E-37 | 35 | Y |
| Chloroflexi bacterium isolate RBG_16_54_18 | Chloroflexi | Rifle well D04 (Rifle, CO, USA) | OGO29581.1 | 8.E-40 | 34 | Y |
| Chloroflexi bacterium isolate SpSt-271 | Chloroflexi | hot spring sediment (British Columbia, Canada) | HDY04783.1 | 4.E-35 | 35 | Y |
| Chloroflexi bacterium isolate SpSt-326 | Chloroflexi | hot spring sediment (British Columbia, Canada) | HDV36418.1 | 2.E-36 | 35 | Y |
| Chloroflexi bacterium isolate SpSt-327 | Chloroflexi | hot spring sediment (British Columbia, Canada) | HEG74209.1 | 5.E-38 | 36 | Y |
| Chloroflexi bacterium isolate SpSt-422 | Chloroflexi | hot spring sediment (British Columbia, Canada) | HFN11568.1 | 5.E-34 | 32 | Y |
| Chloroflexi bacterium isolate SpSt-438 | Chloroflexi | hot spring sediment (British Columbia, Canada) | HFM60814.1 | 7.E-33 | 34 | Y |
| Chloroflexi bacterium isolate SpSt-474 | Chloroflexi | hot spring sediment (British Columbia, Canada) | HGU24517.1 | 2.E-37 | 35 | Y |
| Chloroflexi bacterium isolate SpSt-552 | Chloroflexi | hot spring sediment (British Columbia, Canada) | HGQ02489.1 | 2.E-31 | 32 | Y |
| Chloroflexi bacterium isolate SpSt-583 | Chloroflexi | hot spring sediment (British Columbia, Canada) | HGT18656.1 | 3.E-33 | 34 | Y |
| Chloroflexi bacterium isolate SpSt-600 | Chloroflexi | hot spring sediment (Tengchong, China) | HGO05051.1 | 8.E-34 | 32 | Y |
| Chloroflexi bacterium isolate SpSt-660 | Chloroflexi | hot spring sediment (Tengchong, China) | HGL63430.1 | 3.E-33 | 32 | Y |
| Chloroflexi bacterium isolate SpSt-996 | Chloroflexi | mud, sediment (Yellowstone National Park, USA) | HFU29806.1 | 2.E-33 | 34 | Y |
| Chloroflexi bacterium isolate UBA9854 | Chloroflexi | anaerobic digester | HAI37108.1 | 2.E-26 | 31 | Y |
| Chloroflexi bacterium isolate UTCFX4 | Chloroflexi | anaerobic digester filtrate from the belt filter press of CVWRF wastewater treatment plant (Salt Lake City, UT, USA) | OQY79586.1 | 2.E-23 | 31 | Y |
| Cloacibacillus evryensis strain DSM 19522 | Synergistetes | sewage sludge (Évry, France) | WP_034444262.1 | 6.E-15 | 26 | N |
| Cloacibacillus porcorum strain 105753 | Synergistetes | pig feces | NMF17534.1 | 3.E-14 | 26 | N |
| Cloacibacillus porcorum strain CL-84 | Synergistetes | pig cecal mucosa | ANZ46032.1 | 8.E-12 | 24 | N |
| Cloacibacillus sp. strain An23 | Synergistetes | red junglefowl cecum | OOU93278.1 | 9.E-16 | 27 | N |
| Deferribacter desulfuricans strain SSM1 | Deferribacteres | deep-sea hydrothermal vent (Suiyo Seamount, Japan) | BAI81501.1 | 9.E-05 | 24 | Y |
| Dehalococcoidia bacterium isolate Baikal-deep-G109 | Chloroflexi | Lake Baikal (Russia) | MSQ14392.1 | 3.E-13 | 26 | N |
| Deltaproteobacteria bacterium isolate 1MN72D_58_314 | Proteobacteria | groundwater (Munich, Germany) | TDB36245.1 | 2.E-12 | 28 | Y |
| Deltaproteobacteria bacterium isolate AS06rmzACSIP_532 | Proteobacteria | anaerobic digestion of organic wastes under variable temperature conditions and feedstocks | NLV24503.1 | 2.E-13 | 30 | Y |
| Deltaproteobacteria bacterium isolate AS06rmzACSIP_589 | Proteobacteria | anaerobic digestion of organic wastes under variable temperature conditions and feedstocks | NLX52857.1 | 3.E-38 | 35 | Y |
| Deltaproteobacteria bacterium isolate AS4AgIBPMA_32 | Proteobacteria | anaerobic digestion of organic wastes under variable temperature conditions and feedstocks | NMC97429.1 | 3.E-38 | 35 | Y |
| Deltaproteobacteria bacterium isolate B3_G2 | Proteobacteria | deep-sea hydrothermal vent sediment (Guaymas Basin, Mexico) | RLC06739.1 | 3.E-17 | 28 | Y |
| Deltaproteobacteria bacterium isolate B13_G4 | Proteobacteria | deep-sea hydrothermal vent sediment (Guaymas Basin, Mexico) | RLC22860.1 | 2.E-24 | 30 | Y |
| Deltaproteobacteria bacterium isolate B17_G16 | Proteobacteria | deep-sea hydrothermal vent sediment (Guaymas Basin, Mexico) | RLB37252.1 | 7.E-12 | 26 | Y |
| Deltaproteobacteria bacterium isolate B23_G16 | Proteobacteria | deep-sea hydrothermal vent sediment (Guaymas Basin, Mexico) | RLC12904.1 | 9.E-19 | 31 | Y |
| Deltaproteobacteria bacterium isolate B46_G9 | Proteobacteria | deep-sea hydrothermal vent sediment (Guaymas Basin, Mexico) | RLB18191.1 | 1.E-19 | 29 | Y |
| Deltaproteobacteria bacterium isolate B125_G9 | Proteobacteria | deep-sea hydrothermal vent sediment (Guaymas Basin, Mexico) | RLC29483.1 | 6.E-19 | 26 | Y |
| Deltaproteobacteria bacterium isolate B133_G9 | Proteobacteria | deep-sea hydrothermal vent sediment (Guaymas Basin, Mexico) | RLC20422.1 | 1.E-24 | 32 | N |
| Deltaproteobacteria bacterium isolate B144_G9 | Proteobacteria | deep-sea hydrothermal vent sediment (Guaymas Basin, Mexico) | RLC25857.1 | 9.E-18 | 28 | Y |
| Deltaproteobacteria bacterium isolate CG_4_8_14_3_um_filter_51_11 | Proteobacteria | aquifer (Crystal Geyser, UT, USA) | PIX18886.1 | 2.E-18 | 27 | N |
| Deltaproteobacteria bacterium isolate CG03_land_8_20_14_0_80_45_14 | Proteobacteria | aquifer (Crystal Geyser, UT, USA) | PIV21324.1 | 1.E-22 | 27 | Y |
| Deltaproteobacteria bacterium isolate DOLJRAL78_50_10 | Proteobacteria | gingival sulcus from 5 year old male Dolphin_J | PIE73735.1 | 7.E-21 | 31 | Y |
| Deltaproteobacteria bacterium isolate DOLJRAL78_53_22 | Proteobacteria | gingival sulcus from 5 year old male Dolphin_J | PIE70991.1 | 4.E-07 | 25 | N |
| Deltaproteobacteria bacterium isolate HGW-Deltaproteobacteria-1 | Proteobacteria | groundwater (Horonobe, Japan) | PKN88290.1 | 3.E-30 | 32 | Y |
| Deltaproteobacteria bacterium isolate HGW-Deltaproteobacteria-5 | Proteobacteria | groundwater (Horonobe, Japan) | PKN10657.1 | 2.E-34 | 33 | Y |
| Deltaproteobacteria bacterium isolate HGW-Deltaproteobacteria-6 | Proteobacteria | groundwater (Horonobe, Japan) | PKN18563.1 | 3.E-36 | 35 | Y |
| Deltaproteobacteria bacterium isolate HGW-Deltaproteobacteria-11 | Proteobacteria | groundwater (Horonobe, Japan) | PKN59704.1 | 2.E-33 | 33 | Y |
| Deltaproteobacteria bacterium isolate HGW-Deltaproteobacteria-19 | Proteobacteria | groundwater (Horonobe, Japan) | PKN34680.1 | 1.E-21 | 32 | Y |
| Deltaproteobacteria bacterium isolate INTA.AUR.1003 | Proteobacteria | first pond of the AUR serial dairy industry stabilization pond system (Vila, Argentina) | NCC25723.1 | 1.E-14 | 28 | Y |
| Deltaproteobacteria bacterium isolate J040 | Proteobacteria | iron-rich hot spring (Shikinejima Island, Japan) | RMG60923.1 | 1.E-23 | 31 | N |
| Deltaproteobacteria bacterium isolate MAG 48 | Proteobacteria | hydrothermal vent (Pacific Ocean) | RTZ96695.1 | 3.E-15 | 27 | Y |
| Deltaproteobacteria bacterium isolate RIFOXYC2_FULL_48_10 | Proteobacteria | Rifle well CD01 (Rifle, CO, USA) | OGQ92707.1 | 3.E-22 | 29 | Y |
| Deltaproteobacteria bacterium isolate SpSt-510 | Proteobacteria | hot spring sediment (British Columbia, Canada) | HFH11051.1 | 2.E-20 | 30 | Y |
| Deltaproteobacteria bacterium isolate SpSt-772 | Proteobacteria | mud, sediment (Yellowstone National Park, USA) | HGV81120.1 | 2.E-16 | 27 | Y |
| Deltaproteobacteria bacterium isolate SpSt-871 | Proteobacteria | mud, sediment (Yellowstone National Park, USA) | HGZ73194.1 | 7.E-16 | 27 | Y |
| Deltaproteobacteria bacterium isolate SpSt-1015 | Proteobacteria | mud, sediment (Yellowstone National Park, USA) | HHH87581.1 | 9.E-20 | 30 | Y |
| Deltaproteobacteria bacterium isolate UBA10529 | Proteobacteria | oil sands and coal beds | HAR96709.1 | 2.E-15 | 28 | Y |
| Desulfamplus magnetovallimortis strain BW-1 | Proteobacteria | brackish spring (Death Valley, CA, USA) | SLM31755.1 | 7.E-20 | 25 | Y |
| Desulfatibacillum aliphaticivorans strain AK-01 | Proteobacteria | Arthur Kill, NJ/NY waterway (USA) | ACL02166.1 | 5.E-19 | 33 | N |
| Desulfatibacillum aliphaticivorans strain DSM 15576 | Proteobacteria | hydrocarbon-polluted marine sediment (Gulf of Fos, France) | WP_051327074.1 | 5.E-20 | 34 | N |
| Desulfatibacillum alkenivorans strain DSM 16219 | Proteobacteria | oil-polluted sediment of operation station of ballast water and tank water cleaning (Fos Harbor, France) | WP_083611154.1 | 1.E-17 | 32 | N |
| Desulfatiglans anilini strain DSM 4660 | Proteobacteria | marine sediment (North Sea coast, Germany) | WP_028322618.1 | 1.E-22 | 31 | N |
| Desulfatitalea sp. isolate Site_C24 | Proteobacteria | beach sand (Middle Park Beach, Australia) | NNK02546.1 | 1.E-12 | 28 | Y |
| Desulfatitalea tepidiphila strain S28bF | Proteobacteria | marine sediment (Tokyo Bay, Japan) | WP_076750459.1 | 2.E-23 | 30 | Y |
| Desulfobacter curvatus strain DSM 3379 | Proteobacteria | marine mud (Venice, Italy) | WP_020586554.1 | 2.E-20 | 30 | Y |
| Desulfobacter hydrogenophilus strain AcRS1 | Proteobacteria | marine mud (Venice, Italy) | QBH12555.1 | 8.E-15 | 27 | Y |
| Desulfobacter postgatei strain 2ac9 | Proteobacteria | anaerobic sediment of brackish water ditch (Jadebusen, Germany) | EIM62640.1 | 1.E-21 | 30 | Y |
| Desulfobacter sp. isolate UBA12168 | Proteobacteria | sediment | HBT90065.1 | 2.E-21 | 30 | Y |
| Desulfobacter vibrioformis strain DSM 8776 | Proteobacteria | water-oil separation system on oil platform (North Sea, Norway) | WP_035235263.1 | 4.E-18 | 29 | Y |
| Desulfobacteraceae bacterium isolate 4484_190.3 | Proteobacteria | deep-sea hydrothermal vent sediment (Guaymas Basin, Mexico) | OPX36574.1 | 8.E-19 | 28 | Y |
| Desulfobacteraceae bacterium isolate 4572_88 | Proteobacteria | deep-sea hydrothermal vent sediment (Guaymas Basin, Mexico) | OQY55368.1 | 3.E-20 | 31 | N |
| Desulfobacteraceae bacterium isolate 4572_123 | Proteobacteria | deep-sea hydrothermal vent sediment (Guaymas Basin, Mexico) | OQY04121.1 | 5.E-26 | 30 | Y |
| Desulfobacteraceae bacterium isolate CG2_30_51_40 | Proteobacteria | groundwater (Crystal Geyser, UT, USA) | OIP43152.1 | 2.E-18 | 27 | N |
| Desulfobacteraceae bacterium isolate DS_bin_10 | Proteobacteria | beach sand (Middle Park Beach, Australia) | NNG02102.1 | 2.E-12 | 25 | Y |
| Desulfobacteraceae bacterium isolate Eth-SRB1 | Proteobacteria | anaerobic ethane-degrading enrichment culture inoculated with marine sediment (Gulf of Mexico) | RZB29647.1 | 5.E-18 | 27 | Y |
| Desulfobacteraceae bacterium isolate Eth-SRB2 | Proteobacteria | anaerobic ethane-degrading enrichment culture inoculated with marine sediment (Gulf of Mexico) | RZB36540.1 | 3.E-25 | 30 | Y |
| Desulfobacteraceae bacterium isolate HyVt-13 | Proteobacteria | deep-sea hydrothermal vent sediment (Guaymas Basin, Mexico) | HDI59857.1 | 1.E-10 | 29 | N |
| Desulfobacteraceae bacterium isolate HyVt-208 | Proteobacteria | deep-sea hydrothermal vent sediment (Guaymas Basin, Mexico) | HDL07956.1 | 2.E-22 | 28 | Y |
| Desulfobacteraceae bacterium isolate IS3 | Proteobacteria | microbial mat in sulfidic groundwater-fed sinkhole (Lake Huron, USA) | OQX21718.1 | 1.E-23 | 27 | N |
| Desulfobacteraceae bacterium isolate IS3 | Proteobacteria | microbial mat in sulfidic groundwater-fed sinkhole (Lake Huron, USA) | OQW99265.1 | 2.E-19 | 30 | Y |
| Desulfobacteraceae bacterium isolate IS3 | Proteobacteria | microbial mat in sulfidic groundwater-fed sinkhole (Lake Huron, USA) | OQX26173.1 | 2.E-18 | 30 | Y |
| Desulfobacteraceae bacterium isolate Madre2 | Proteobacteria | bioplastic (PHA) biofilm from water-sediment interface of coastal lagoon (Upper Laguna Madre, TX, USA) | THB81596.1 | 3.E-17 | 27 | Y |
| Desulfobacteraceae bacterium isolate SURF_4 | Proteobacteria | deeply circulating subsurface aquifer fluids (Lead, SD, USA) | RJQ76849.1 | 7.E-11 | 26 | Y |
| Desulfobacteraceae bacterium isolate SURF_4 | Proteobacteria | deeply circulating subsurface aquifer fluids (Lead, SD, USA) | RJQ65780.1 | 2.E-05 | 21 | N |
| Desulfobacteraceae bacterium isolate SURF_7 | Proteobacteria | deeply circulating subsurface aquifer fluids (Lead, SD, USA) | RJP79134.1 | 6.E-21 | 29 | Y |
| Desulfobacteraceae bacterium isolate UBA8212 | Proteobacteria | seawater | HCY85389.1 | 9.E-14 | 28 | Y |
| Desulfobacteraceae bacterium isolate UBA8400 | Proteobacteria | microbial mat (Lake Huron, MI, USA) | HAO21021.1 | 5.E-27 | 27 | N |
| Desulfobacterales bacterium isolate DOLJRAL78_48_6 | Proteobacteria | gingival sulcus from 5 year old male Dolphin_J | PIE60952.1 | 7.E-18 | 28 | Y |
| Desulfobacterales bacterium isolate GLR701 | Proteobacteria | Glendhu Ridge methane seep (Pacific Ocean) | NOQ19417.1 | 9.E-24 | 30 | N |
| Desulfobacterales bacterium isolate HyVt-57 | Proteobacteria | deep-sea hydrothermal vent sediment (Guaymas Basin, Mexico) | HGY11719.1 | 1.E-17 | 28 | Y |
| Desulfobacterales bacterium isolate HyVt-57 | Proteobacteria | deep-sea hydrothermal vent sediment (Guaymas Basin, Mexico) | HGY11834.1 | 4.E-15 | 28 | Y |
| Desulfobacterales bacterium isolate HyVt-59 | Proteobacteria | deep-sea hydrothermal vent sediment (Guaymas Basin, Mexico) | HHC24326.1 | 1.E-23 | 32 | Y |
| Desulfobacterales bacterium isolate S7086C20 | Proteobacteria | methane seep sediment (Pacific Ocean) | OEU45016.1 | 4.E-24 | 31 | Y |
| Desulfobacterales bacterium isolate Site_B14 | Proteobacteria | beach sand (Middle Park Beach, Australia) | NNK84523.1 | 2.E-23 | 29 | Y |
| Desulfobacterales bacterium isolate SpSt-563 | Proteobacteria | hot spring sediment (British Columbia, Canada) | HGP66397.1 | 6.E-19 | 30 | Y |
| Desulfobacterales bacterium isolate UWMA-0216 | Proteobacteria | Mid-Cayman Rise vent fluids (Atlantic Ocean) | HID59813.1 | 2.E-15 | 26 | Y |
| Desulfobacterium autotrophicum strain HRM2 | Proteobacteria | marine sediment (Mediterranean Sea) | ACN17620.1 | 3.E-19 | 30 | Y |
| Desulfobacterium vacuolatum strain DSM 3385 | Proteobacteria | marine mud (Venice, Italy) | SMC85487.1 | 7.E-17 | 29 | Y |
| Desulfobacula phenolica strain DSM 3384 | Proteobacteria | marine mud (Venice, Italy) | SDU45768.1 | 3.E-16 | 26 | Y |
| Desulfobacula sp. isolate GWF2_41_7 | Proteobacteria | Rifle well CD01 (Rifle, CO, USA) | OGR15363.1 | 5.E-19 | 27 | Y |
| Desulfobacula sp. isolate SM1_1_2 | Proteobacteria | stromatolite mat (Schoenmakerskop, South Africa) | NJM02770.1 | 1.E-17 | 28 | N |
| Desulfobacula toluolica strain Tol2 | Proteobacteria | marine mud (Woods Hole, MA, USA) | CCK82146.1 | 2.E-15 | 26 | Y |
| Desulfobulbaceae bacterium isolate DB1 | Proteobacteria | anode biofilm in microbial fuel cells | OKY74567.1 | 6.E-05 | 27 | Y |
| Desulfobulbus oralis strain HOT041/ORNL | Proteobacteria | human subgingival sample | AVD70372.1 | 2.E-57 | 37 | Y |
| Desulfobulbus oralis strain HOT041/ORNL | Proteobacteria | human subgingival sample | AVD71509.1 | 1.E-61 | 40 | Y |
| Desulfococcus multivorans strain DSM 2059 | Proteobacteria | sewage digester (Göttingen, Germany) | EPR35993.1 | 2.E-19 | 32 | Y |
| Desulfococcus sp. isolate 4484_241 | Proteobacteria | deep-sea hydrothermal vent sediment (Guaymas Basin, Mexico) | OQX63297.1 | 3.E-20 | 29 | N |
| Desulfococcus sp. isolate 4484_242 | Proteobacteria | deep-sea hydrothermal vent sediment (Guaymas Basin, Mexico) | OQX61744.1 | 2.E-16 | 28 | N |
| Desulfoluna spongiiphila strain AA1 | Proteobacteria | marine sponge Aplysina aerophoba (Mediterranean Sea) | SCX79058.1 | 2.E-06 | 27 | N |
| Desulfoluna spongiiphila strain DBB | Proteobacteria | marine intertidal sediment (L'Escala, Spain) | VVS90461.1 | 8.E-06 | 26 | N |
| Desulfomonile tiedjei isolate SpSt-769 | Proteobacteria | mud, sediment (Yellowstone National Park, USA) | HGH61094.1 | 5.E-28 | 32 | Y |
| Desulfomonile tiedjei strain DSM 6799 | Proteobacteria | sewage sludge (Adrian, MI, USA) | AFM25414.1 | 1.E-36 | 35 | Y |
| Desulfonatronospira thiodismutans strain ASO3-1 | Proteobacteria | sediment from a highly alkaline saline soda lake on the Kulunda Steppe (Altai, Russia) | EFI33801.1 | 1.E-22 | 33 | Y |
| Desulfonema ishimotonii strain Tokyo 01 | Proteobacteria | marine sediment (Tokyo Bay, Japan) | GBC64034.1 | 3.E-21 | 30 | N |
| Desulfosarcina alkanivorans strain PL12 | Proteobacteria | oil spill of shallow marine sediment (Shuaiba, Kuwait) | BBO72287.1 | 2.E-19 | 30 | Y |
| Desulfosarcina cetonica strain JCM 12296 | Proteobacteria | flooded oil stratum of the Apsheron peninsula (Azerbaijan) | WP_054702774.1 | 2.E-13 | 28 | N |
| Desulfosarcina ovata subsp. sediminis strain 28bB2T | Proteobacteria | tidal flat sediment (Tokyo Bay, Japan) | BBO80496.1 | 3.E-11 | 27 | Y |
| Desulfosarcina sp. strain Bu55 | Proteobacteria | sediment (Guaymas Basin, Mexico) | WP_027353905.1 | 9.E-15 | 25 | Y |
| Desulfosarcina widdellii strain PP31 | Proteobacteria | oil spill of shallow marine sediment (Shuaiba, Kuwait) | BBO74481.1 | 8.E-15 | 26 | Y |
| Desulfospira joergensenii strain DSM 10085 | Proteobacteria | marine surface sediment below sea grass (Arcachon Bay, France) | WP_022666032.1 | 2.E-19 | 28 | Y |
| Desulfotignum balticum strain DSM 7044 | Proteobacteria | marine mud (Saxild, Denmark) | WP_024336727.1 | 6.E-23 | 32 | Y |
| Desulfotignum phosphitoxidans strain DSM 13687 | Proteobacteria | marine sediment (Venice, Italy) | EMS79033.1 | 3.E-23 | 32 | Y |
| Desulfovibrio sp. strain An276 | Proteobacteria | red junglefowl cecum | OOU52973.1 | 3.E-60 | 35 | Y |
| Desulfovibrionaceae bacterium isolate CIM:MAG 1040 | Proteobacteria | human feces | PWM70546.1 | 1.E-65 | 41 | Y |
| Desulfurispirillum indicum strain S5 | Chrysiogenetes | river sediment (Chennai, India) | ADU66823.1 | 1.E-04 | 26 | Y |
| Desulfurivibrio sp. isolate SURF_16 | Proteobacteria | deeply circulating subsurface aquifer fluids (Lead, SD, USA) | RJX31147.1 | 8.E-11 | 28 | Y |
| Desulfuromonadales bacterium isolate C00003093 | Proteobacteria | sediment from site of active methane seepage at Hydrate Ridge South (Pacific Ocean) | OEU75150.1 | 2.E-18 | 28 | Y |
| Fibrobacter sp. isolate AS06rmzACSIP_199 | Fibrobacteres | anaerobic digestion of organic wastes under variable temperature conditions and feedstocks | NLG16780.1 | 3.E-22 | 29 | Y |
| Fibrobacter sp. isolate AS06rmzACSIP_529 | Fibrobacteres | anaerobic digestion of organic wastes under variable temperature conditions and feedstocks | NLD91845.1 | 1.E-21 | 27 | N |
| Fibrobacter sp. isolate AS06rmzACSIP_543 | Fibrobacteres | anaerobic digestion of organic wastes under variable temperature conditions and feedstocks | NLE01532.1 | 7.E-21 | 30 | Y |
| Fibrobacter sp. isolate AS09scLD_71 | Fibrobacteres | anaerobic digestion of organic wastes under variable temperature conditions and feedstocks | NLP03213.1 | 2.E-22 | 30 | Y |
| Fibrobacter sp. isolate AS22ysBPME_152 | Fibrobacteres | anaerobic digestion of organic wastes under variable temperature conditions and feedstocks | NLL13864.1 | 3.E-20 | 27 | Y |
| Fibrobacteres bacterium isolate UBA10882 | Fibrobacteres | wetland surface sediment | HAJ80221.1 | 6.E-20 | 30 | Y |
| Firmicutes bacterium isolate AS23ysBPME_266 | Firmicutes | anaerobic digestion of organic wastes under variable temperature conditions and feedstocks | NMB11274.1 | 4.E-06 | 25 | N |
| Flexilinea flocculi strain TC1 | Chloroflexi | methanogenic granular sludge (Belgium) | GAP39404.1 | 5.E-43 | 34 | Y |
| Fretibacterium sp. isolate AS21ysBPME_7 | Synergistetes | anaerobic digestion of organic wastes under variable temperature conditions and feedstocks | NLL37650.1 | 1.E-16 | 32 | N |
| Geovibrio thiophilus strain DSM 11263 | Deferribacteres | drainage ditch (Konstanz, Germany) | QAR33452.1 | 4.E-08 | 28 | Y |
| Ignavibacteriae bacterium isolate B3_G1 | Chlorobi | deep-sea hydrothermal vent sediment (Guaymas Basin, Mexico) | RKY90554.1 | 4.E-22 | 29 | Y |
| Lentimonas sp. CC4 | Verrucomicrobia | seawater (Nahant, MA, USA) | CAA6679502.1 | 1.E-15 | 30 | Y |
| Lentimonas sp. CC10 | Verrucomicrobia | seawater (Nahant, MA, USA) | CAA6691588.1 | 3.E-15 | 29 | Y |
| Lentisphaerae bacterium isolate AS06rmzACSIP_375 | Lentisphaerae | anaerobic digestion of organic wastes under variable temperature conditions and feedstocks | NLE68485.1 | 5.E-15 | 29 | Y |
| Lentisphaerae bacterium isolate RIFOXYA12_64_32 | Lentisphaerae | Rifle well CD01 (Rifle, CO, USA) | OGV62430.1 | 2.E-28 | 33 | Y |
| Leptolinea sp. isolate AS06rmzACSIP_266 | Chloroflexi | anaerobic digestion of organic wastes under variable temperature conditions and feedstocks | NLF50870.1 | 2.E-41 | 35 | Y |
| Leptolinea sp. isolate AS27yjCOA_114 | Chloroflexi | anaerobic digestion of organic wastes under variable temperature conditions and feedstocks | NMB53094.1 | 2.E-36 | 34 | Y |
| Leptolinea tardivitalis strain YMTK-2 | Chloroflexi | wastewater from a factory producing shochu (Yamagawa Beach, Japan) | KPL70500.1 | 2.E-38 | 33 | Y |
| Levilinea saccharolytica strain KIBI-1 | Chloroflexi | sludge granules from sugar-processing plant (Niigata, Japan) | KPL75587.1 | 3.E-35 | 35 | Y |
| Nitrolancea hollandica strain Lb | Chloroflexi | nitrifying bioreactor (Rotterdam, Netherlands) | CCF84771.1 | 8.E-06 | 24 | N |
| Nitrospira moscoviensis strain NSP M-1 | Nitrospirae | eroded iron pipe (Moscow, Russia) | ALA58029.1 | 2.E-16 | 29 | Y |
| Olavius algarvensis Delta 1 endosymbiont | Proteobacteria | shallow marine sediment (Italy) | CAB1077770.1 | 1.E-21 | 29 | Y |
| Olavius sp. associated proteobacterium Delta 1 | Proteobacteria | shallow marine sediment (Italy) | CAB1061238.1 | 6.E-20 | 28 | Y |
| Ornatilinea apprima strain P3M-1 | Chloroflexi | microbial mat formed in wooden bath filled with hot water from a 2775 m-deep well (Siberia, Russia) | KPL78704.1 | 1.E-20 | 33 | Y |
| Pelodictyon luteolum strain DSM 273 | Chlorobi | meromictic lake (Norway) | ABB23996.1 | 2.E-15 | 29 | Y |
| Pelodictyon phaeoclathratiforme strain BU-1 | Chlorobi | monimolimnion of Buchensee (Germany) | ACF43535.1 | 1.E-22 | 32 | Y |
| Pelolinea submarina strain DSM 23923 | Chloroflexi | marine subsurface sediment (Pacific Ocean) | BBB47806.1 | 2.E-35 | 32 | Y |
| Planctomycetes bacterium isolate GSL.Bin20 | Planctomycetes | microbial mat (Bridger Bay, UT, USA) | NBB94753.1 | 5.E-22 | 30 | Y |
| Prosthecochloris aestuarii isolate SpSt-1181 | Chlorobi | hot spring sediment (Wilbur Springs, CA, USA) | HED30548.1 | 2.E-28 | 36 | Y |
| Prosthecochloris aestuarii strain DSM 271 | Chlorobi | hydrogen sulfide containing mud of brackish lagoon (Lake Sasyk-Sivash, Russia) | ACF46288.1 | 2.E-20 | 30 | N |
| Prosthecochloris marina strain V1 | Chlorobi | coastal area of the South China Sea | PWW83239.1 | 5.E-23 | 31 | N |
| Prosthecochloris sp. isolate C10 | Chlorobi | seawater lake chemocline (Dragon's Eye Lake, Croatia) | NEX14508.1 | 3.E-17 | 28 | N |
| Prosthecochloris sp. strain CIB 2401 | Chlorobi | coastal lagoon (Mallorca, Spain) | ANT65030.1 | 2.E-26 | 34 | Y |
| Prosthecochloris sp. strain GSB1 | Chlorobi | deep-sea hydrothermal vent (Pacific Ocean) | ASQ91644.1 | 2.E-26 | 33 | N |
| Prosthecochloris sp. strain HL-130-GSB | Chlorobi | phototrophic microbial mat (Hot Lake, WA, USA) | ARM30833.1 | 5.E-28 | 36 | Y |
| Prosthecochloris sp. strain ZM_2 | Chlorobi | water from chemokine of meromictic lakes Green cape (Karelia, Russia) | RNA64855.1 | 8.E-24 | 34 | Y |
| Prosthecochloris vibrioformis strain DSM 260 | Chlorobi | rivermouth | TNJ36903.1 | 9.E-27 | 34 | Y |
| Proteobacteria bacterium isolate DOLZORAL124_55_4 | Proteobacteria | gingival sulcus from 29 year old lactating female Dolphin_Z | PID40410.1 | 2.E-14 | 28 | Y |
| Proteobacteria bacterium isolate SpSt-1152 | Proteobacteria | hot spring sediment (Wilbur Springs, CA, USA) | HEC99630.1 | 2.E-15 | 28 | Y |
| Pseudodesulfovibrio sp. strain S3 | Proteobacteria | Black Sea water (Bulgaria) | RWU02434.1 | 2.E-06 | 26 | Y |
| Smithella sp. isolate AS06rmzACSIP_551 | Proteobacteria | anaerobic digestion of organic wastes under variable temperature conditions and feedstocks | NLD79966.1 | 2.E-35 | 33 | Y |
| Smithella sp. isolate AS17jrsBPGN_1 | Proteobacteria | anaerobic digestion of organic wastes under variable temperature conditions and feedstocks | NLA41131.1 | 1.E-32 | 33 | Y |
| Smithella sp. isolate AS4AgIBPMA_11 | Proteobacteria | anaerobic digestion of organic wastes under variable temperature conditions and feedstocks | NMC92510.1 | 5.E-35 | 34 | Y |
| Smithella sp. isolate D17 | Proteobacteria | produced water from an oilfield (Medicine Hat, Alberta, Canada) | KFZ45035.1 | 2.E-36 | 34 | Y |
| Smithella sp. isolate F21 | Proteobacteria | oil sands tailings pond (Medicine Hat, Alberta, Canada) | KFN39154.1 | 7.E-36 | 34 | Y |
| Smithella sp. isolate SCADC | Proteobacteria | mature fine tailings from oil sands tailings pond (Medicine Hat, Alberta, Canada) | KFO68937.1 | 3.E-37 | 35 | Y |
| Smithella sp. isolate SDB | Proteobacteria | sediment (San Diego Bay, CA, USA) | KQC09736.1 | 6.E-33 | 33 | Y |
| Smithella sp. isolate UBA10513 | Proteobacteria | oil sands and coal beds | HBI47682.1 | 1.E-38 | 34 | Y |
| Sphaerobacter thermophilus strain DSM 20745 | Chloroflexi | thermal treated municipal sewage sludge (Muenchen-Grosslappen, Germany) | ACZ37929.1 | 9.E-07 | 28 | N |
| Sphaerochaeta dissipatitropha strain GLS2 | Spirochaetes | arctic permafrost (Russia) | SMP46469.1 | 1.E-21 | 31 | Y |
| Sphaerochaeta globosa strain Buddy | Spirochaetes | marine hot spring (Shiashkoten Island, Russia) | ADY12711.1 | 8.E-22 | 30 | Y |
| Sphaerochaeta halotolerans strain 4-11 | Spirochaetes | Production water from oilfield (Vostochno-Anziskoe oilfield, Russia) | RFU94749.1 | 1.E-17 | 31 | Y |
| Sphaerochaeta pleomorpha strain Grapes | Spirochaetes | Red Cedar River (Okemos, MI, USA) | AEV28959.1 | 1.E-18 | 30 | Y |
| Sphaerochaeta sp. isolate MAG5 | Spirochaetes | Red-pigmented microbial biofilm developed on the inner wall of a bioreactor (Jena, Germany) | TAH57281.1 | 2.E-17 | 30 | Y |
| Sphaerochaeta sp. isolate UBA8956 | Spirochaetes | oil sands and coal beds | HAF86241.1 | 4.E-12 | 26 | Y |
| Sphaerochaeta sp. isolate UBA9948 | Spirochaetes | oil sands and coal beds | HAP57835.1 | 9.E-20 | 31 | Y |
| Sphaerochaeta sp. isolate UBA11053 | Spirochaetes | bioreactor | HCU30833.1 | 2.E-17 | 30 | Y |
| Sphaerochaeta sp. isolate UBA11175 | Spirochaetes | oil sands and coal beds | HBO36282.1 | 7.E-20 | 31 | Y |
| Spirochaetales bacterium isolate AS06rmzACSIP_457 | Spirochaetes | anaerobic digestion of organic wastes under variable temperature conditions and feedstocks | NLE15601.1 | 7.E-20 | 30 | Y |
| Spirochaetales bacterium isolate AS07pgkLD_77 | Spirochaetes | anaerobic digestion of organic wastes under variable temperature conditions and feedstocks | NLY07479.1 | 1.E-14 | 30 | Y |
| Spirochaetales bacterium isolate AS08sgBPME_324 | Spirochaetes | anaerobic digestion of organic wastes under variable temperature conditions and feedstocks | HHT81641.1 | 2.E-17 | 29 | Y |
| Spirochaetales bacterium isolate AS10tIH2TH_379 | Spirochaetes | anaerobic digestion of organic wastes under variable temperature conditions and feedstocks | NLA97632.1 | 2.E-20 | 30 | Y |
| Spirochaetales bacterium isolate AS22ysBPME_10 | Spirochaetes | anaerobic digestion of organic wastes under variable temperature conditions and feedstocks | NLL25178.1 | 4.E-17 | 29 | Y |
| Spirochaetales bacterium isolate AS22ysBPME_249 | Spirochaetes | anaerobic digestion of organic wastes under variable temperature conditions and feedstocks | NMA22686.1 | 3.E-15 | 30 | Y |
| Spirochaetales bacterium isolate AS23ysBPME_118 | Spirochaetes | anaerobic digestion of organic wastes under variable temperature conditions and feedstocks | NLK13860.1 | 8.E-18 | 29 | Y |
| Spirochaetales bacterium isolate AS23ysBPME_120 | Spirochaetes | anaerobic digestion of organic wastes under variable temperature conditions and feedstocks | NLK06812.1 | 1.E-15 | 29 | Y |
| Spirochaetes bacterium isolate ADurb.Bin315 | Spirochaetes | anaerobic digester (Urbana, IL, USA) | OQA43369.1 | 4.E-16 | 28 | Y |
| Spirochaetes bacterium strain RBG_16_49_21 | Spirochaetes | Rifle well D04 (Rifle, CO, USA) | OHD69784.1 | 2.E-12 | 27 | N |
| Spirochaetia bacterium isolate INTA.CYC.017 | Spirochaetes | first pond of the CYC serial dairy industry stabilization pond system (San Carlos Sur, Argentina) | NCC13313.1 | 4.E-17 | 28 | Y |
| Spirochaetia bacterium isolate UBA12135 | Spirochaetes | aquifer (Colorado River, CO, USA) | HBE03855.1 | 4.E-05 | 24 | Y |
| Synergistaceae bacterium isolate AS07pgkLD_87 | Synergistetes | anaerobic digestion of organic wastes under variable temperature conditions and feedstocks | NLV82264.1 | 5.E-11 | 26 | N |
| Synergistes sp. strain 3_1_syn1 | Synergistetes | human gut | EHL68284.1 | 7.E-15 | 26 | N |
| Synergistetes bacterium isolate HGW-Synergistetes-1 | Synergistetes | groundwater (Horonobe, Japan) | PKL03800.1 | 4.E-08 | 25 | N |
| Syntrophaceae bacterium isolate UBA10514 | Proteobacteria | oil sands and coal beds | HBH86494.1 | 7.E-19 | 30 | Y |
| Syntrophaceae bacterium isolate UBA11395 | Proteobacteria | oil sands and coal beds | HBJ74262.1 | 3.E-38 | 35 | Y |
| Syntrophobacter sp. strain SbD1 | Proteobacteria | peat soil (Weissenstadter Forst-Nord, Germany) | SPF48485.1 | 2.E-24 | 34 | Y |
| Syntrophus sp. isolate UBA8958 | Proteobacteria | oil sands and coal beds | HAI26831.1 | 1.E-21 | 32 | Y |
| Syntrophus sp. isolate UBA10520 | Proteobacteria | oil sands and coal beds | HAR97668.1 | 2.E-18 | 31 | Y |
| Thermaerobacter sp. strain PB12/4term | Firmicutes | Sediment (Lake Baikal, Russia) | QIA27848.1 | 4.E-04 | 28 | N |
| uncultured Desulfatiglans sp. isolate IK1 | Proteobacteria | asphalt lake (Pitch Lake, Trinidad and Tobago) | VBB43204.1 | 3.E-19 | 32 | N |
| uncultured Desulfobacteraceae bacterium isolate CR-1 | Proteobacteria | ectosymbiote of a magnetic protist of Mediterranean sea | VEN73776.1 | 2.E-15 | 28 | Y |
| Uncultured Desulfobacterium sp. | Proteobacteria | contaminated aquifer (Stuttgart, Germany) | CBX28553.1 | 2.E-24 | 30 | N |
| Verrucomicrobia bacterium isolate DP16D_bin.41 | Verrucomicrobia | groundwater (Oak Ridge, TN, USA) | TAN38092.1 | 4.E-22 | 29 | Y |

**Table S2.**
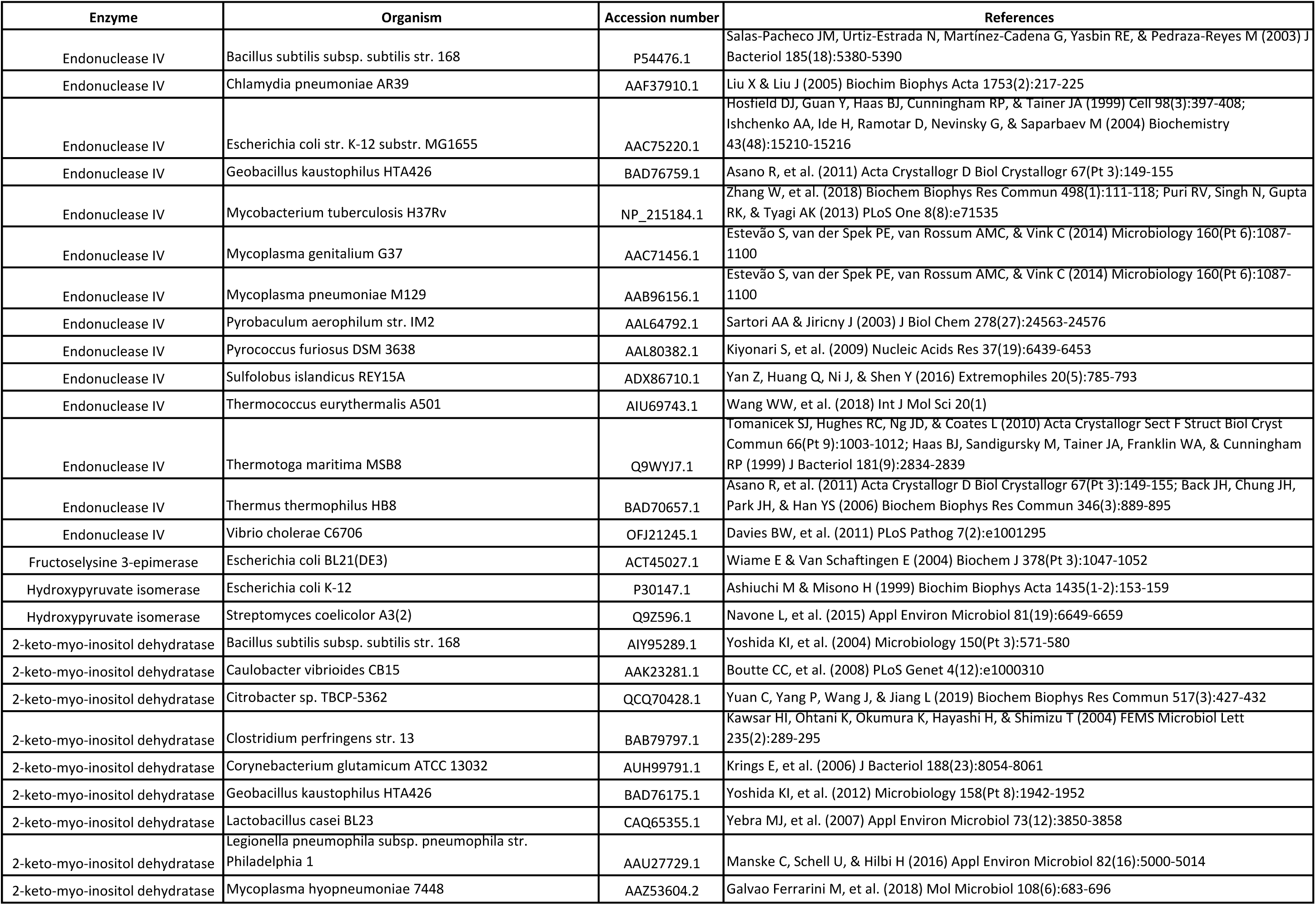
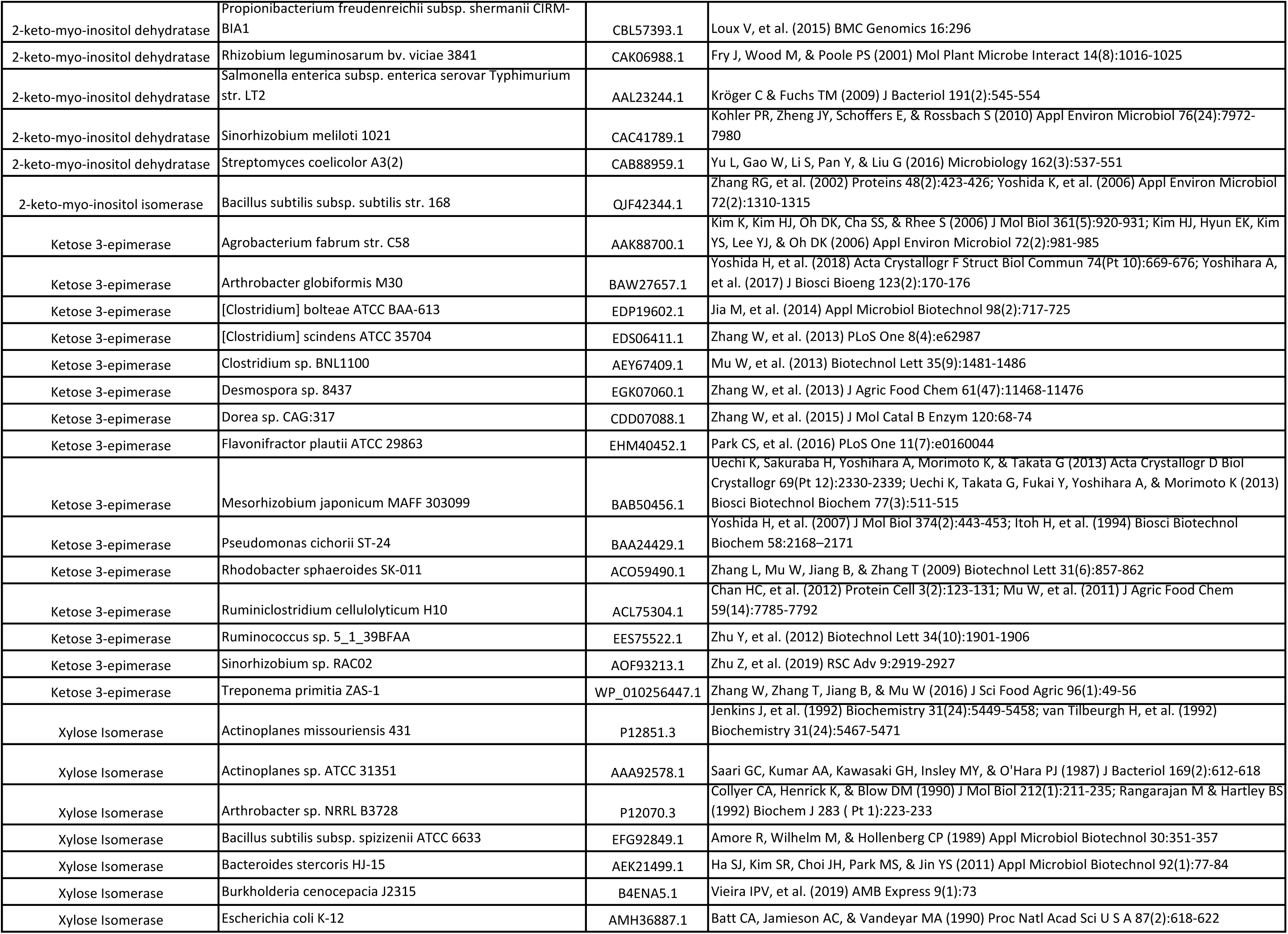
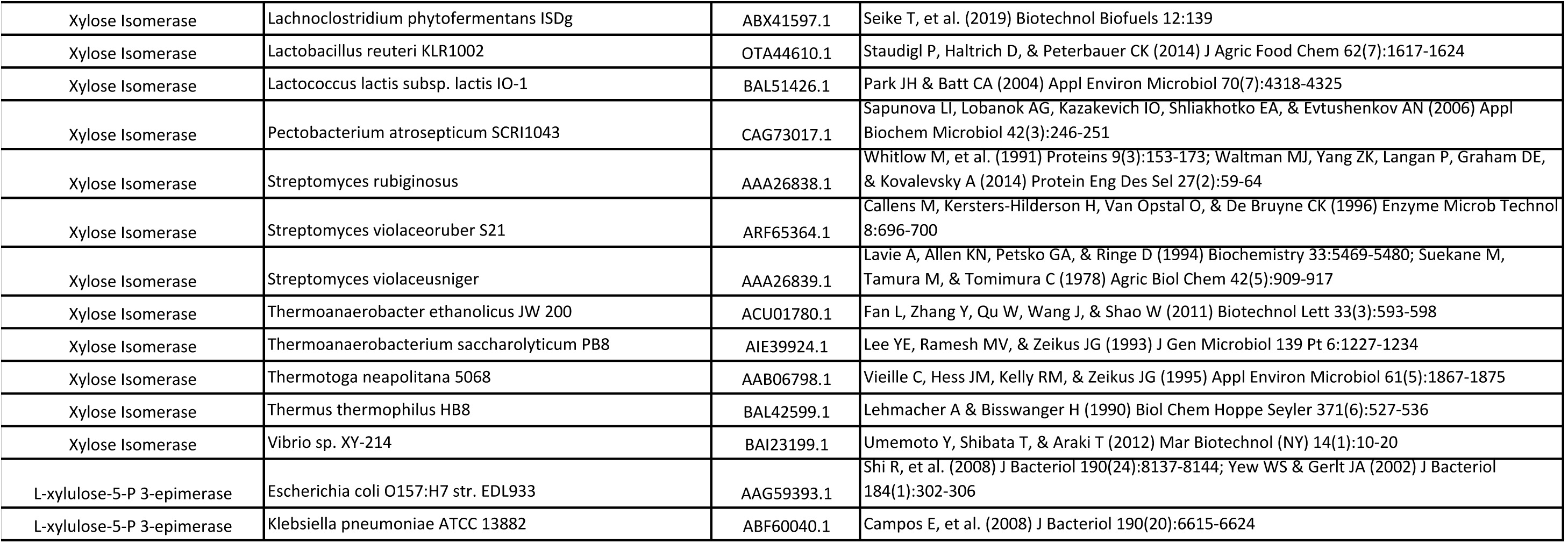
Biochemically and structurally characterized members of AP endonuclease 2 superfamily

